# Epstein-Barr Virus Latent Membrane Protein 1 Suppresses Ferroptosis via Pentose Phosphate Pathway and Glutathione Metabolism

**DOI:** 10.64898/2026.03.09.710628

**Authors:** Eric M. Burton, Bidisha Mitra, Rui Guo, John M. Asara, Benjamin E. Gewurz

## Abstract

Epstein-Barr virus (EBV) is associated with 200,000 cancers per year, including Burkitt lymphoma and post-transplant lymphomas. We previously reported that EBV latency oncogene programs dynamically remodel infected B cell metabolism and sensitivity to induction of ferroptosis, a programmed cell death pathway driven by lipid reactive oxygen species. However, much has remained unknown about how EBV remodels key redox defense pathways in support of infected B cell proliferation. Here, we identify EBV latent membrane protein 1 (LMP1), a key viral oncogene necessary for B cell immortalization and which mimics aspects of CD40 signaling, drives resistance to ferroptosis induction by erastin, a small molecule that blocks cystine uptake. LMP1 expression was sufficient to protect Burkitt cells from erastin ferroptosis induction. Mechanistically, signaling from the LMP1 TES2/CTAR2 region drove this phenotype, which was not shared by CD40 signaling, revealing that LMP1 evolved independent redox defense roles. Metabolomic analysis highlighted key LMP1 and TES2 signaling roles in support of antioxidant cysteine and glutathione levels. TES2 signaling supported cystine uptake, glutathione and NADPH pools in newly infected peripheral blood B cells. We identified PFKFB4, a host enzyme that shunts glucose into the pentose phosphate pathway to support NADPH production, as a major TES2 metabolic target. PFKFB4 knockdown increased EBV-transformed lymphoblastoid cell line lipid ROS levels, decreased glutathione and strongly sensitized them to ferroptosis induction by erastin treatment. PFKFB4 was also necessary for LMP1-mediated Burkitt B cell ferroptosis resistance. Collectively, these results identify PFKFB4 as a key host cell EBV metabolism remodeling target critical for infected B cell redox defense.

## Introduction

The gamma-herpesvirus Epstein-Barr virus (EBV) persistently infects most adults worldwide. EBV was the first human tumor virus discovered(1)(2) and is now associated with a wide range of human malignancies. EBV is a major driver of Burkitt lymphoma, Hodgkin lymphoma, post-transplant lymphoproliferative diseases, central nervous system lymphoma, nasopharyngeal and gastric carcinoma(3–6). EBV is also increasingly associated with autoimmunity and may serve as a major viral trigger for multiple sclerosis and systemic lupus erythematosus(7–11).

EBV uses a series of viral oncoproteins to drive the outgrowth and differentiation of newly infected cells, different subsets of which are expressed in EBV-associated cancers. EBV transforms primary human B cells into continuously growing, immortalized lymphoblastoid cell lines (LCLs) in cell culture. LCLs are an excellent model system for EBV-driven immunoblastic lymphomas of immunosuppressed hosts, in particular PTLD. EBV uses a series of epigenetically regulated latency programs to transform primary B-cells into LCLs, w(12)hich express eight EBV-encoded oncoproteins and non-coding RNAs. These include six Epstein-Barr nuclear antigens (EBNA) and two latent membrane proteins (LMP). LMP1 mimics aspects of B cell co-receptor CD40 signaling(13–15), whereas LMP2A mimics aspects of B cell immunoglobulin receptor signaling(16, 17).

LMP1 is comprised of a short N-terminal cytoplasmic tail and six transmembrane domains that drive ligand-independent signaling from a 200 residue C-terminal cytoplasmic tail(18–21). Genetic studies identified two LMP1 signaling regions within the C-terminal tail that are critical for EBV-mediated conversion of primary human B cells into LCLs(22, 23). These were termed transformation effector site (TES) 1 and 2 and are also referred to as C-terminal activating regions 1 and 2 (CTAR1/2). TES1 signaling is required for initiation of B-cell outgrowth, whereas TES2 signaling is required for later stages of EBV-driven B-cell immortalization(24–28). Likewise, signaling by both LMP1 C-terminal tail domains is critical for LCL survival(29). Interestingly however, B cells infected by EBV with point mutations that abrogate TES2 signaling can be cultured for long periods of time on irradiated fibroblast feeders(25), a major function of which is to support cysteine metabolism by converting extracellular cystine (the oxidized form of cysteine) into cysteine, which can facilitate B cell uptake(30, 31).

To support rapid B-cell outgrowth, EBV strongly remodels newly infected B-cell metabolism. EBV highly induces lipid metabolism pathways as it converts resting B cells into blasts that undergo Burkitt-like hyperproliferation between days 3-7 post-infection and express highly elevated levels of the metabolism master regulator MYC oncogene(32, 33). As LMP1 and LMP2A levels increase, infected cells convert from Burkitt-like to lymphoblastoid B cell physiology. However, much remained to be learned about how EBV, and LMP1 signaling in particular, alters infected B-cell immunometabolism.

Elevated lipid metabolism can produce lipid reactive oxygen species (ROS) byproducts. If not detoxified, lipid ROS can cause catastrophic damage to host cell membranes and trigger the ferroptosis programmed cell death pathway(34, 35). A key redox defense pathway utilizes the host enzyme GPX4, which uses reduced glutathione as a cofactor to detoxify lipid ROS species(36). Glutathione is a peptide synthesized from cysteine, glutamine and glycine, with cysteine the rate-limiting amino acid. Cystine uptake, reduction to cysteine and incorporation into GPX4 therefore play major roles in defense against ferroptosis induction(37). We found that Burkitt cells with the EBV latency I program, in which EBNA1 is the only EBV-encoded protein expressed, are exquisitely sensitive to ferroptosis induction by inhibition of cystine uptake or by GPX4 blockade(38), mirroring the important observation that Burkitt cells undergo oxidative stress when cultured at low density(39). In fact, GPX4 was discovered in a screen for factors that protect Burkitt B cells from oxidative stress(40). By contrast, we found that LCLs are resistant to ferroptosis induction

EBV dynamically sensitizes newly infected primary human B cells to ferroptosis induction(41). Whereas B cells undergoing Burkitt-like hyperproliferation are highly sensitive to ferroptosis induction, as cells convert to lymphoblasts with increased LMP1 expression, transforming cells become progressively resistant to ferroptosis induction, particularly by inhibition of cystine uptake. Whereas newly infected cells undergoing Burkitt-like hyper-proliferation are exquisitely sensitive to ferroptosis induction, LCL-like exhibit resistance to ferroptosis induction. However, the mechanisms underlying these phenotypes, and their potential relationship to LMP1, have remained unknown.

Here, we identified that LMP1 plays a major role in remodeling B-cell redox defense pathways. Genetic and metabolomic approaches highlighted that LMP1 TES2 signaling support cystine uptake, NADPH production for its conversion to cysteine and for expansion of reduced glutathione pools in support of ferroptosis evasion. Remarkably, LMP1 TES2 signaling was found to be sufficient to protect Burkitt cells from ferroptosis induction by the cystine uptake inhibitor erastin. The host metabolic enzyme PFKFB4 was identified as a key LMP1 TES2 target, whose expression was necessary for LCL defense against erastin ferroptosis induction, identifying it as a therapeutic target.

## Results

### LMP1 promotes Burkitt B cell ferroptosis resistance

Based open our prior observation that EBV progressively induces ferroptosis resistance in newly infected cells, particularly to SLC7A11 blockade, and the correlation of this phenotype with timepoints at which LMP1 and LMP2A expression levels increase(41), we hypothesized that LMP1 and/or LMP2A signaling drives B cell ferroptosis resistance. We therefore investigated whether expression of LMP1 or LMP2A, each of which signal constitutively in a ligand-independent manner, was sufficient to confer resistance to ferroptosis induction in Daudi Burkitt lymphoma cells. To do so, we engineered Daudi cells with doxycycline-inducible LMP1 or LMP2A expression. We then tested ferroptosis induction by the SLC7A11 inhibitor erastin or by the GPX4 inhibitor ML-210(42, 43) in cells mock induced or induced for LMP1 or LMP2A expression for 24 hours. As expected, mock induced Daudi Burkitt cells were exquisitely sensitive to ferroptosis induction by erastin or ML-210, and this was rescuable by treatment with the lipophilic antioxidant Fer-1(44). Interestingly, LMP1 but not LMP2A expression strongly conferred resistance to ferroptosis induction by erastin treatment but not by ML-210 (**Fig. 1A-B, S1A-B**). LMP1 expression also strongly suppressed erastin-induced lipid ROS production, as judged by FACS analysis of cells stained by BODIPY 581/591 C-11, a membrane-localizing fluorescent lipid peroxidation dye (**Fig. 1C-D**). 8-point dose titration highlighted that LMP1 expression markedly increased the resistance to erastin but not ML-210 in both EBV+ Daudi and EBV-Akata Burkitt cells, suggesting that LMP1 itself, rather than LMP1 induction of another EBV gene, conferred ferroptosis resistance (**Fig. 1E-H**). Taken together, these data suggest that LMP1 blocks ferroptosis induction at a step in the pathway prior to GPX4-mediated detoxification of lipid ROS.

**Figure 1:**
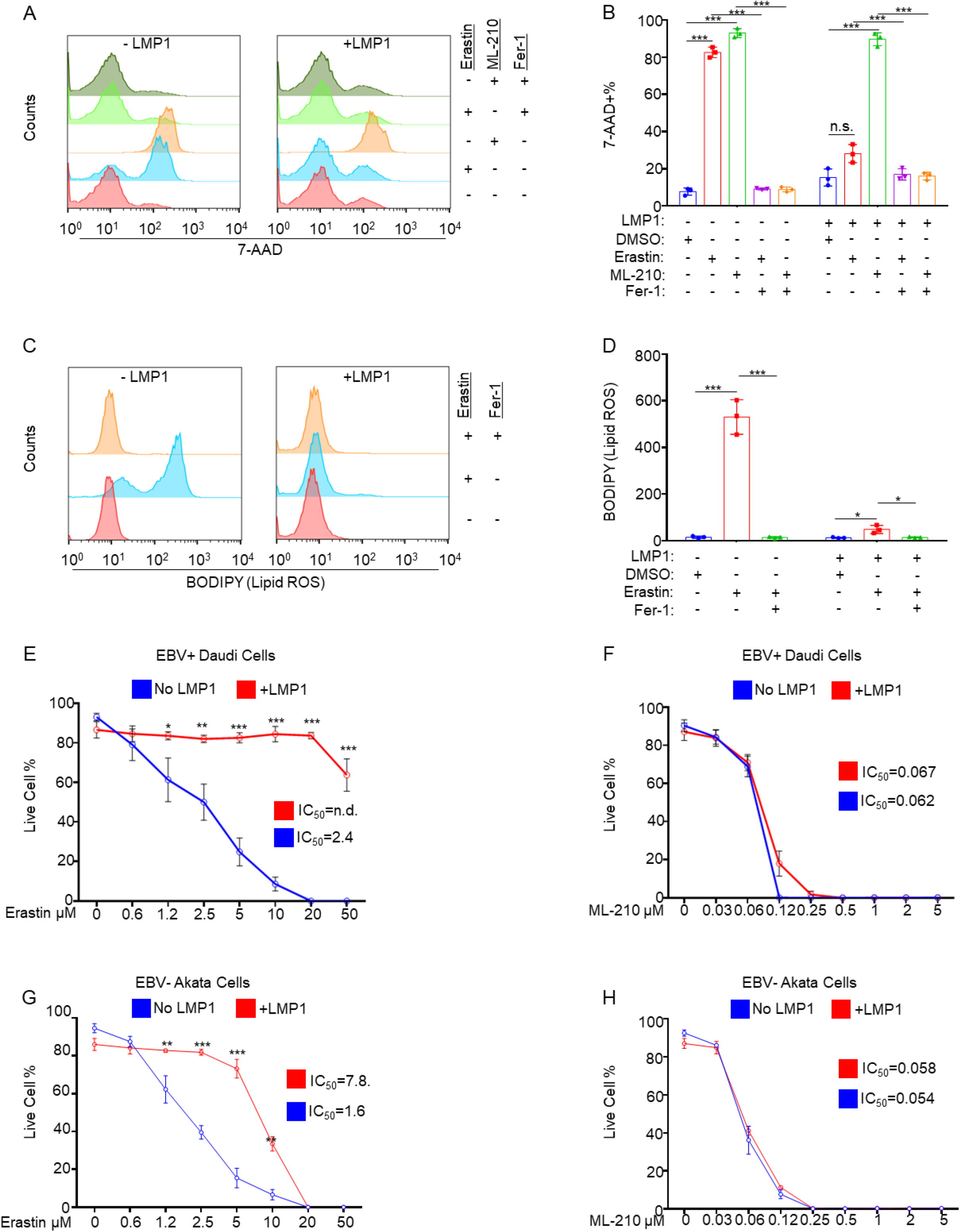
LMP1 prevents erastin-induced ferroptosis. (A) Analysis of LMP1 effects on Burkitt cell ferroptosis induction by erastin or ML-210. Representative FACS analysis of 7-AAD uptake by Daudi cells mock induced or induced for LMP1 expression for 24 hours by 250ng/mL doxycycline and then treated with DMSO, 10µM erastin (SLC7A11 inhibitor), 0.1µM ML-210 (GPX4 inhibitor) and/or 2.5µM Ferrostatin-1 (Fer1) for 48 hours. (B) Mean ± SD percentages of 7-AAD+ cells from n=3 replicates as in (A). (C) Analysis of LMP1 effects on lipid peroxidation induction in Burkitt cells. Representative FACS analysis of lipid ROS levels of Daudi cells mock induced or induced for LMP1 expression for 24 hours by dox and then treated with DMSO vehicle, 10µM erastin and/or 2.5µM Fer-1 for 18 hours, followed by BODIPY C-11 staining. (D) Mean ± SD MFI of BODIPY C-11 membrane lipid ROS staining from n=3 replicates as in (C). (E) IC_50_ dose curve analysis of LMP1-expressing Burkitt cells in response to erastin. Live cell percentage analysis of Daudi cells mock-induced or induced for LMP1 expression for 24 hours and then treated with the indicated doses of erastin for 24 hours before viability analysis via CellTiter Glo. (F) IC_50_ dose curve analysis of LMP1-expressing Burkitt cells in response to erastin. Live cell percentage analysis of Daudi cells mock induced or induced for LMP1 expression for 24 hours and then treated with the indicated doses of ML-210 for 24 hours before viability analysis via CellTiter Glo. (G) IC_50_ dose curve analysis of LMP1-expressing Burkitt cells in response to erastin. Live cell percentage analysis of Akata-EBV-negative cells mock induced or induced for LMP1 expression for 24 hours and then treated with the indicated doses of erastin for 24 hours before viability analysis via CellTiter Glo. (H) IC_50_ dose curve analysis of LMP1-expressing Burkitt cells in response to erastin. Live cell percentage analysis of Akata-EBV-negative cells mock induced or induced for LMP1 expression for 24 hours and then treated with the indicated doses of ML-210 for 24 hours before viability analysis via CellTiter Glo. P-values were determined by one-sided Fisher’s exact test. * p<0.05, **p<0.005, ***p<0.0005.

### LMP1 transformation effector site 2 signaling promotes ferroptosis resistance

LMP1 mimics CD40 signaling to activate oncogenic signaling pathways, including NF-κB, MAP kinase, PI3 kinase, protein kinase C and IRF7 (**Fig. 2A**) (45–50). Therefore, to test whether CD40 signaling likewise induces erastin resistance, we stimulated Daudi cells with multimerized CD40 ligand (CD40L). As LMP1-mediated canonical and non-canonical NF-κB activation are required for lymphoblastoid B-cell survival, we also treated cells in parallel with the SMAC mimetic birinapant(51) and with tumor necrosis factor α (TNFα) to preferentially activate non-canonical versus canonical NF-κB signaling, respectively (**Fig. S2**). Interestingly, although each agonist activated cellular signaling, only SMAC mimetic treatment conferred erastin resistance. However, SMAC mimetic treatment significantly impaired Daudi cell fitness (**Figure 2B, S3A-B**).

**Figure 2.**
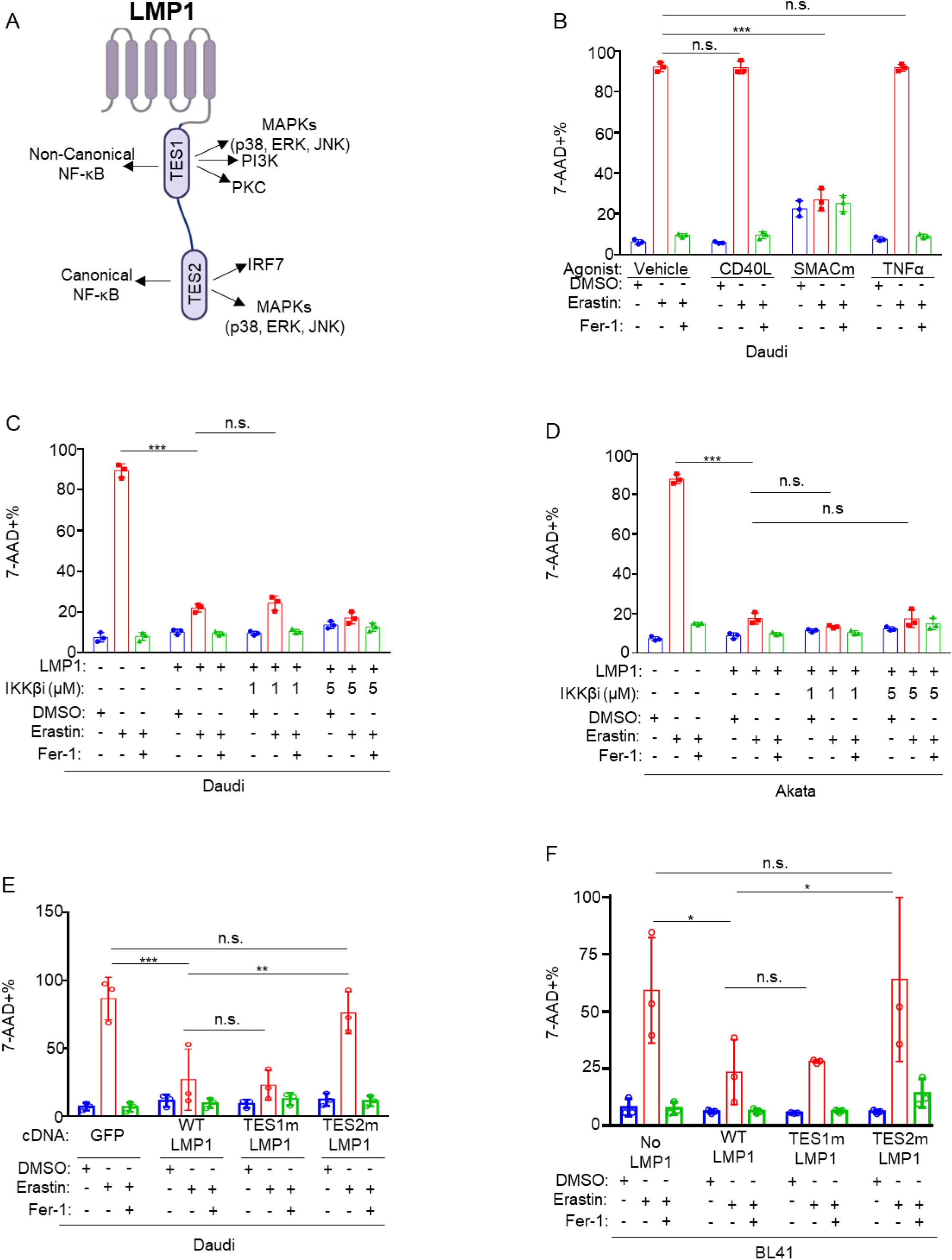
LMP1 protection from erastin ferroptosis induction is dependent on TES2 signaling but not dependent on canonical NF-κB. (A) Model of pathways activated by LMP1 transformation effector sites (TES) 1 and 2. Created in BioRender. Burton, E. (2026) https://BioRender.com/225hby0 (B) Analysis of immunoreceptor signaling effects on Burkitt cell erastin ferroptosis induction. FACS %7AAD+ mean + SD values from n=3 independent replicates of Daudi cells stimulated with DMSO control, MEGACD40L (50ng/mL), SMAC mimetic birinapant (20µM) or TNFα (10ng/mL). After 24 hours, cells were treated with DMSO, 10μM erastin and/or 2.5μM Fer-1. Viability was assessed 48 hours later. (C) Analysis of canonical NF-κB roles on LMP1-mediated ferroptosis protection. FACS %7AAD+ mean + SD values from n=3 independent replicates of Daudi cells mock induced or dox induced for LMP1 along with DMSO or the indicated doses of IKKβ inhibitor IKK-2 VIII for 24 hours. Cells were then treated with DMSO vehicle, 10µM erastin and/or 5µM Fer-1 for 48 hours quantitation of live cell numbers. (D) Analysis of canonical NF-κB roles in LMP1-mediated ferroptosis protection. FACS %7AAD+ mean + SD values from n=3 independent replicates of Akata cells mock induced or dox-induced for LMP1 expression for 24 hours along with DMSO vehicle or the indicated doses of IKKβ inhibitor IKK-2 VIII for 24 hours. Cells were then treated with DMSO, 10µM erastin and/or 2.5µM Fer-1 for 48 hours and cell death was quantitated by FACS measurement of 7-AAD uptake. (E) Analysis of LMP1 TES domain contribution to ferroptosis resistance. FACS %7AAD+ mean + SD values from n=3 independent replicates of Daudi cells mock induced or dox-induced for expression of control GFP, wild-type LMP1 or point mutant LMP1 abrogated for TES1 or TES2 signaling. 24 hours later, cells were treated with DMSO vehicle, 10µM erastin and/or 2.5µM Fer-1 for 48 hours and cell death was quantitated by FACS measurement of 7-AAD uptake. (F) Analysis of LMP1 TES domain contribution to ferroptosis resistance. FACS %7AAD+ mean + SD values from n=3 independent replicates of BL-41 cells mock induced or induced for control GFP, wild-type LMP1 or point mutant LMP1 abrogated for TES1 or TES2 signaling. 24 hours later, cells were treated with DMSO vehicle, 10µM erastin and/or 2.5µM Fer-1 for 48 hours and cell death was quantitated by FACS measurement of 7-AAD uptake. P-values were determined by one-sided Fisher’s exact test. * p<0.05, **p<0.005, ***p<0.0005.

Given the above results, we hypothesized that LMP1 does not induce erastin resistance via canonical NF-κB induction. To test this hypothesis, we treated Daudi cells with an IKKβ inhibitor at two doses, together with LMP1 induction. At both 1 and 5 μM doses, IKKβ inhibition failed to rescue erastin-driven ferroptosis induction, despite blockade of LMP1 target genes TRAF1 and p100 (**Fig. 2C, S3C-D**). Similar results were obtained in EBV-negative Akata Burkitt B cells (**Fig. 2D, S3E-F**). We therefore next hypothesized that LMP1 TES1 signaling is required for erastin resistance. To test this, we induced Daudi cells for conditional expression of control green fluorescence protein (GFP), for wildtype (WT) LMP1, or for well characterized LMP1 point mutants that abrogate signaling from TES1 (TES1m) or from TES2 (TES2m)(26, 52, 53). Expression of WT and TES1m EBV, but not TES2m EBV, induced erastin resistance in both Daudi and BL-41 B cells (**Fig. 2E-F, S4A-D**). These results indicate that LMP1 TES2 signaling confers Burkitt B cell erastin resistance in a manner independent of its activation of canonical NF-κB signaling. However, CRISPR depletion of either TRAF6 or TAK1 which mediate LMP1 TES2 driven canonical NF-κB and MAP kinase activation failed to block erastin resistance driven by conditional LMP1 expression (**Fig. S5-6**). Taken together, these data indicate that TES2 signaling by a distinct pathway, perhaps mediated by TRADD, TRAF2, RIP1 or IRF7, which have all been reported to bind to the TES2/CTAR2 region(47, 54–60).

### LMP1 TES2 signaling regulates EBV-infected B cell glutathione pools

Since LMP1 induced resistance to erastin but not GPX4 blockade, we reasoned that TES2 signaling altered metabolism upstream of GPX4 itself. Importantly, GPX4 uses reduced glutathione (GSH), a tri-peptide comprised of glycine, glutamate and cysteine, as a cofactor to detoxify lipid ROS. Glutathione is the most abundant antioxidant in mammalian cells and cysteine is rate-limiting amino acid for its biosynthesis(36). GSH is converted to oxidized glutathione (GSSG) with GPX4 ROS detoxification.

To gain insights into how EBV latency programs and LMP1 in particular alter metabolites directly relevant to ferroptosis sensitivity versus resistance, we performed liquid chromatography mass spectrometry (LC-MS) analysis of isogenic Mutu Burkitt cells which differ only by EBV latency program. Mutu III expresses all eight EBV latency proteins including LMP1, whereas EBNA1 is the only EBV-encoded protein expressed by Mutu I(61). Interestingly, GSH and to a lesser extent GSSG were significantly more abundant in extracts from Mutu III vs Mutu I cells (**Fig. 3A**). Multiple glycolysis pathway metabolites were more highly detected with latency III, as were metabolites of the pentose phosphate pathway (PPP), including fructose-6-phosphate (F-6-p) and ribose-5-phosphate (R-5-p)(**Fig. 3A-B**). However, PPP metabolite 6-phospho-gluconate was higher in cells with latency I, perhaps suggesting rapid conversion via PPP metabolism to R-5-p in cells with latency III (**Fig. 3A**).

**Figure 3.**
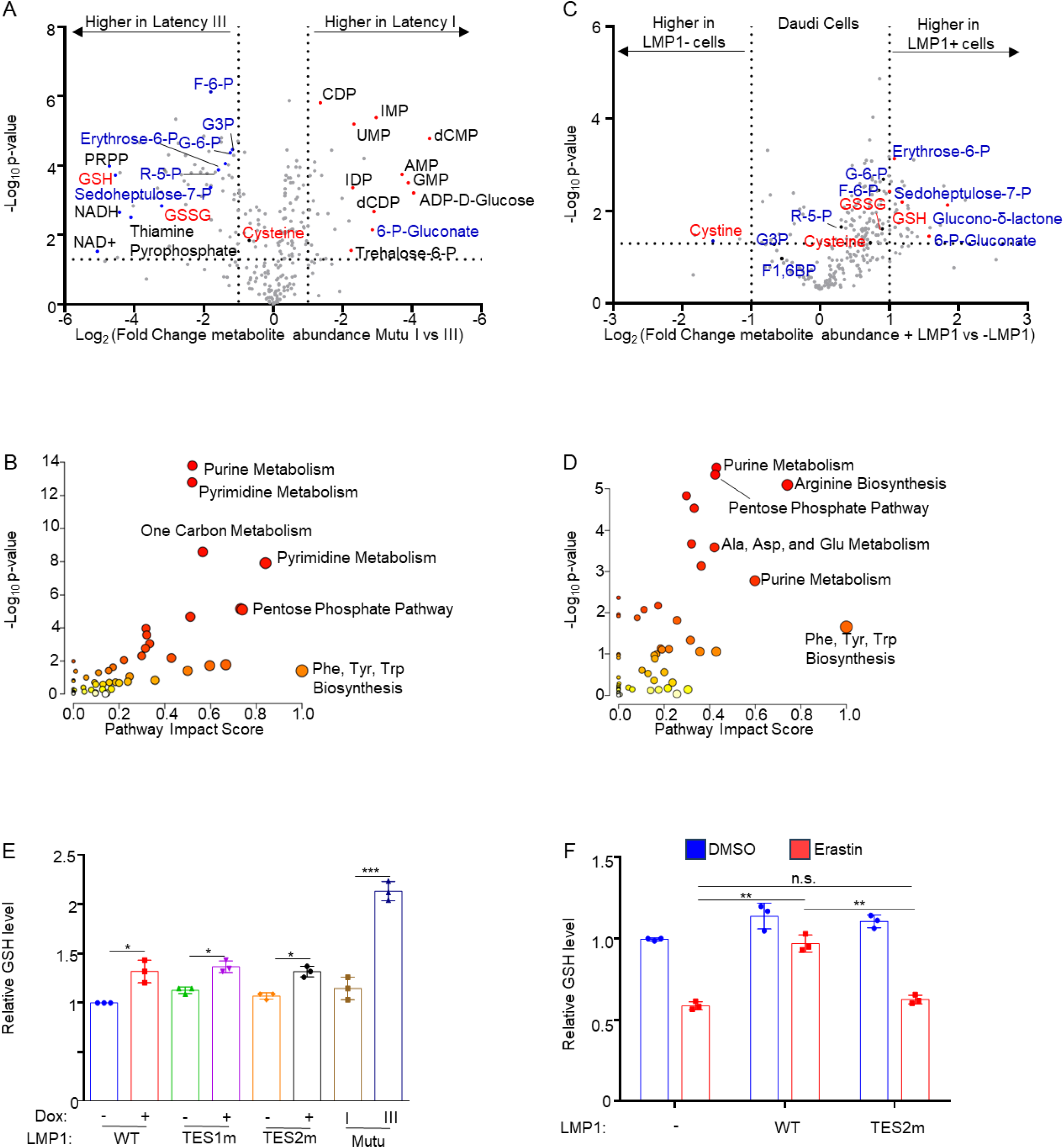
LMP1 signaling supports B cell glutathione pools. (A) Volcano plot of liquid chromatography mass spectrometry (LC-MS) metabolite analysis of n=3 replicates of EBV-infected Mutu cells with latency I vs latency IIl. Positive fold changes indicate higher metabolite concentrations in latency I Mutu cells, while negative fold changes indicate higher metabolite concentrations in latency III Mutu cells. Selected metabolites are indicated. Reduced (GSH) and oxidized (GSSG) glutathione are indicated in red. (B) MetaboAnalyst pathway analysis of metabolites significantly upregulated in latency III versus latency I Mutu cells, as in (A) (C) Volcano plot LC-MS metabolite analysis of n=3 replicates of Daudi cells mock induced or induced for LMP1 expression for 24 hours. Positive fold changes indicate higher metabolite concentrations in LMP1+ vs LMP1-cells. Selected metabolites induced vs. suppressed by LMP1 are indicated. (D) MetaboAnalyst pathway analysis of metabolites significantly upregulated in LMP1-expressing Daudi cells in comparison to mock, as in (C) (E) Daudi cells were mock induced or dox induced for WT, TES1m or TES2m LMP1 expression for 24 hours and then subject to GSH quantitation. Shown are relative GSH levels, normalized by values in mock-induced cells. Mutu I and Mutu III cells were also analyzed. (F) Glutathione colorimetric assay analysis of LMP1-expressing cells in response to cystine uptake inhibition. Daudi Burkitt cells were mock induced or induced for WT or TES2m LMP1 for 24 hours before treatment with DMSO or 10μM erastin for another 18 hours before GSH quantification assay was performed. Shown are relative GSH levels, normalized to values in DMSO treated mock controls. P-values were determined by one-sided Fisher’s exact test. * p<0.05, **p<0.005, ***p<0.0005.

To build upon these results, we next re-analyzed LC-MS data from Daudi cells mock induced or induced for LMP1 expression for 24 hours(62). This analysis highlighted that LMP1 signaling rapidly increased glutathione abundance by ∼4-fold, while cystine levels were depleted by nearly 4-fold (**Fig. 3C-D**). We further investigated latency III and LMP1 effects on B cell glutathione pools via a direct glutathione-detection assay. We again observed that latency III strongly increased GSH levels (**Fig. 3E**). Whereas erastin treatment rapidly depleted GSH from Daudi cells mock-induced for LMP1, LMP1 induction for 24 hours strongly protected cells from erastin-mediated GSH depletion (**Fig. 3F**). By contrast, expression of TES2m LMP1 failed to do so (**Fig. 3F**). Together, these data newly implicate TES2 signaling in B cell cystine and glutathione metabolism for redox defense against ferroptosis induction.

### LMP1 TES2 signaling is required for maintenance of glutathione pools in support of primary human B cell transformation

Classical studies indicated that LMP1 TES1 signaling is necessary for early stages of EBV-driven primary human B cell growth transformation and that LMP1 TES2 signaling subsequently becomes necessary(24, 25, 28). However, it has remained undefined when over the first 6 weeks post-infection that TES2 signaling becomes essential for infected B cell proliferation. To more precisely define this, and to explore a potential linkage between infected B cell dependency on TES2 signaling and redox defense, we used bacterial artificial chromosome (BAC) recombineering to engineer the LMP1 _384_YYD_386_->ID mutation into the EBV genome. Primary human B cells from three independent donors were infected by EBV with WT LMP1, with TES2m _384_YYD_386_->ID. Interestingly, cells infected with TES2m LMP1 grew similarly to those infected with WT LMP1 through approximately day 18 post-infection, at which point the growth curves significantly diverged (**Fig. 4A**).

**Figure 4.**
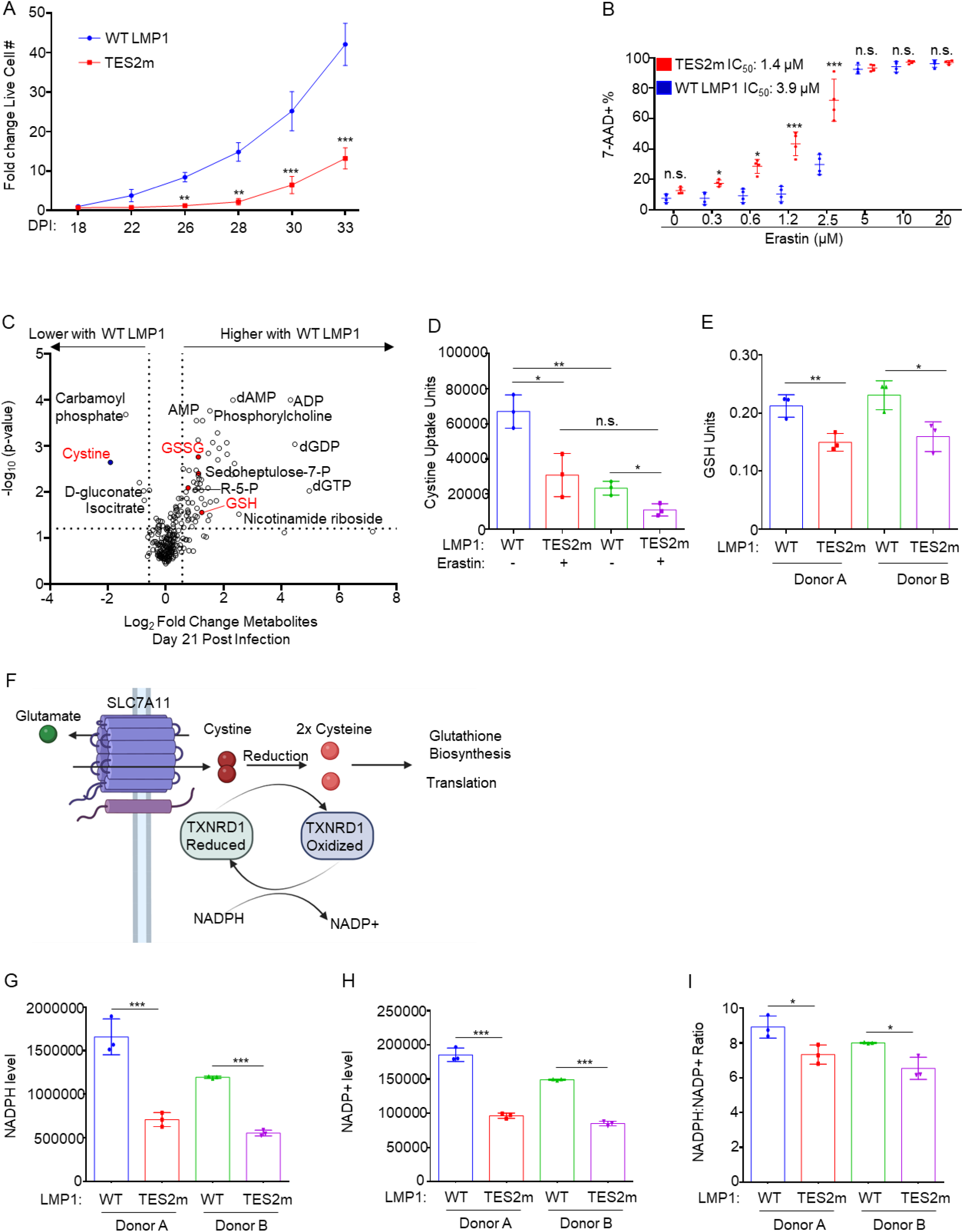
LMP1 TES2 signaling promotes resistance to erastin ferroptosis induction and increases NADPH and GSH pools during B cell immortalization. (A) Growth curve analysis of WT versus TES2m LMP1 EBV infected cells. Peripheral blood B cells were infected with EBV containing WT or TES2m LMP1. After 18 days, cells were re-seeded and growth curve analysis was performed at the indicated days post infection (DPI). Fold change live cell values were calculated relative to numbers at the time of seeding on day 18-post-infection. (B) Analysis of cystine uptake inhibition effects on cells infected by TES2m EBV. Primary B cells from four independent donors were infected with EBV expressing WT or TES2m LMP1. Cells were treated with the indicated erastin doses 21 days post-infection. The %7AAD+ cells was assessed by FACS 48 hours later and IC_50_ of erastin in wild-type versus TES2 mutant infection was calculated. (C) Volcano plot of LC-MS metabolite analysis of n=6 technical replicates averaged from two independent donors. Primary B cells infected with WT or TES2m EBV were analyzed at 21 days post-infection. Positive fold changes indicate higher metabolite concentrations in cells infected by WT EBV, while negative fold changes indicate higher metabolite concentrations in cells infected by TES2m EBV. Cystine, cysteine, and glutathione (GSH) are indicated in red. (D) Effects of LMP1 signaling on cystine uptake. Primary B cells from three independent donors were infected with WT vs TES2m EBV. At 21 days post-infection, cystine uptake assay was performed. As a control, WT and TES2m LMP1 infected cells from one donor were also treated with erastin to demonstrate cystine uptake inhibition. (E) GSH quantification of WT versus TES2m LMP1 infected cells. Primary B cells from three independent donors were infected with WT or TES2m EBV for 21 days, and quantification of GSH levels was performed. (F) Model of cystine metabolism. Cystine is imported via SLC7A11. This oxidized form of cystine is reduced to bio-available cysteine via thioredoxin reductase 1 (TXNRD1) for redox defense and other uses. (G) Effects of WT versus TES2m EBV on infected cell NADPH levels. Primary B cells from two independent donors were infected with WT or TES2m LMP1 EBV. At day 21, NADPH levels were quantitated. (H) Effects of WT versus TES2m EBV on infected cell NADP levels. Primary B cells from two independent donors were infected with WT or TES2m LMP1 EBV. At day 21, NADP levels were quantitated. (I) Effects of WT versus TES2m EBV on NADPH:NADP+ ratios. NADPH:NADP+ ratios from E-F. P-values were determined by one-sided Fisher’s exact test. * p<0.05, **p<0.005, ***p<0.0005.

We next defined sensitivity to erastin ferroptosis induction of cells infected by EBV with WT vs TES2m LMP1 at the point at which the growth curves first strongly diverged. At Day 21 post-infection, cells from 4 independent donors were treated with erastin at serial dilutions between 0-32μM. Whereas B cells infected by EBV with WT LMP1 demonstrated considerable erastin resistance with an IC_50_ of 3.9μM, cells infected with TES2m EBV were significantly more sensitive to erastin induced cell death, with an erastin IC_50_ of 1.4μM (**Fig. 4B**). This data indicates that LMP1 modulates sensitivity to ferroptosis induction at the key timepoint at which transforming B cells become dependent on TES2, likely at the level of cysteine metabolism.

To gain insights into how loss of TES2 signaling alters infected B-cell metabolomes at this key timepoint, we performed LC/MS analysis on B cells from 6 independent donors infected by EBV with WT vs TES2m LMP1. Interestingly, glutathione levels were significantly higher in cells with WT LMP1, whereas cystine was highly depleted, relative to levels observed in cells with TES2m LMP1 (**Fig. 4C, Fig S7 and Supplemental Table 1**). The observation that cystine levels were nearly 4-fold higher in cells infected by EBV with TES2m LMP1 at Day 21 post-infection could be consistent either with increased cystine uptake or with decreased cystine consumption. To distinguish between these, we next directly measured effects of TES2 signaling on primary B cell cystine uptake at day 21 post-infection. Consistent with a key TES2 signaling role in support of cystine uptake, the absence of TES2 signaling strongly impaired cystine uptake at day 21 post-infection (**Fig. 4D**). We validated the significantly lower reduced glutathione levels in cells infected by EBV with TES2M vs WT LMP1 from two additional donors by the GSH abundance assay (**Fig. 4E**). These data indicate that LMP1 TES2 signaling supports both cystine and glutathione metabolism in transforming B cells at several weeks post-infection and support a model in which LMP1 support of glutathione pools supports redox defense. Taken together, these data imply that TES2 mutant LMP1 infection results in reduced cystine uptake, likely as a direct consequence diminished glutathione biosynthesis, given that cysteine is the rate limiting amino acid for GSH production.

Imported cystine is rapidly reduced to cysteine. To do so, cells typically use thioredoxin reductase (TXNRD1), a selenium-containing enzyme that utilizes NADPH as an electron donor to oxidize cystine to cysteine (63) (**Fig. 4F**). We therefore next investigated effects of TES2 mutation on NADP and NADPH pools in primary human B cells at day 21 post-infection. Consistent with key TES2 signaling roles in support of NADPH generation, NADPH levels were significantly lower at day 21 post-infection in B cells infected by TES2m than WT LMP1 EBV (**Fig. 4G**). While NADP levels were also lower with loss of TES2 signaling at this key timepoint, the NADPH/NADP ratio was significantly lower in cells infected by TES2m EBV in either of two independent donors (**Fig. 4H-J**). Taken together, this suggests that LMP1 TES2 maintains NADP/NADPH levels in the cells in support of cystine reduction, glutathione biosynthesis and regeneration of reduced from oxidized glutathione.

### LMP1 TES2 regulates pentose phosphate enzyme PFKFB4 levels during primary B cell infection

To gain further insights into how TES2 signaling regulates infected cell redox defense, we performed RNAseq on primary B cells at day 21 post infection with WT vs. TES2M LMP1 EBV. Notably, mRNA encoding the stress-inducible protein sestrin 2 (SESN2) was highly upregulated in TES2m+ cells (**Fig. 5A, Supplementary Table 2**). SESN2 can be induced by ROS and supports redox homeostasis(64–66). mRNA encoding 6-Phosphofructo-2-Kinase/Fructose-2,6-Biphosphatase 4 (PFKFB4), a major glucose metabolism regulator, was amongst the most highly downmodulated by loss of TES2 signaling at this timepoint (**Fig. 5A**). Interestingly, it is a bifunctional kinase/phosphatase which controls levels of the glycolytic metabolite fructose-2,6-bisphosphate (F2,6BP). PFKFB4 phosphatase activity shunts glucose into the pentose phosphate pathway (PPP) for NADPH production and for *de novo* nucleic acid biosynthesis(67)(**Fig. 5B**). Furthermore, KEGG pathway analysis of mRNAs more highly upregulated by wildtype that LMP1 TES2m EBV highlighted fructose and mannose metabolism as one of the most highly induced pathways (**Fig. 5C**). By contrast, KEGG analysis identified glutathione biosynthesis as one of the more highly induced pathways in cells infected by EBV with TES2m versus wildtype LMP1 (**Fig. 5D**). mRNAs encoding CHAC1, which cleaves and recycles glutathione and OPLAH, which supports glutathione synthesis, perhaps indicative of glutathione depletion at this timepoint of infection by EBV with TES2m LMP1.

**Figure 5.**
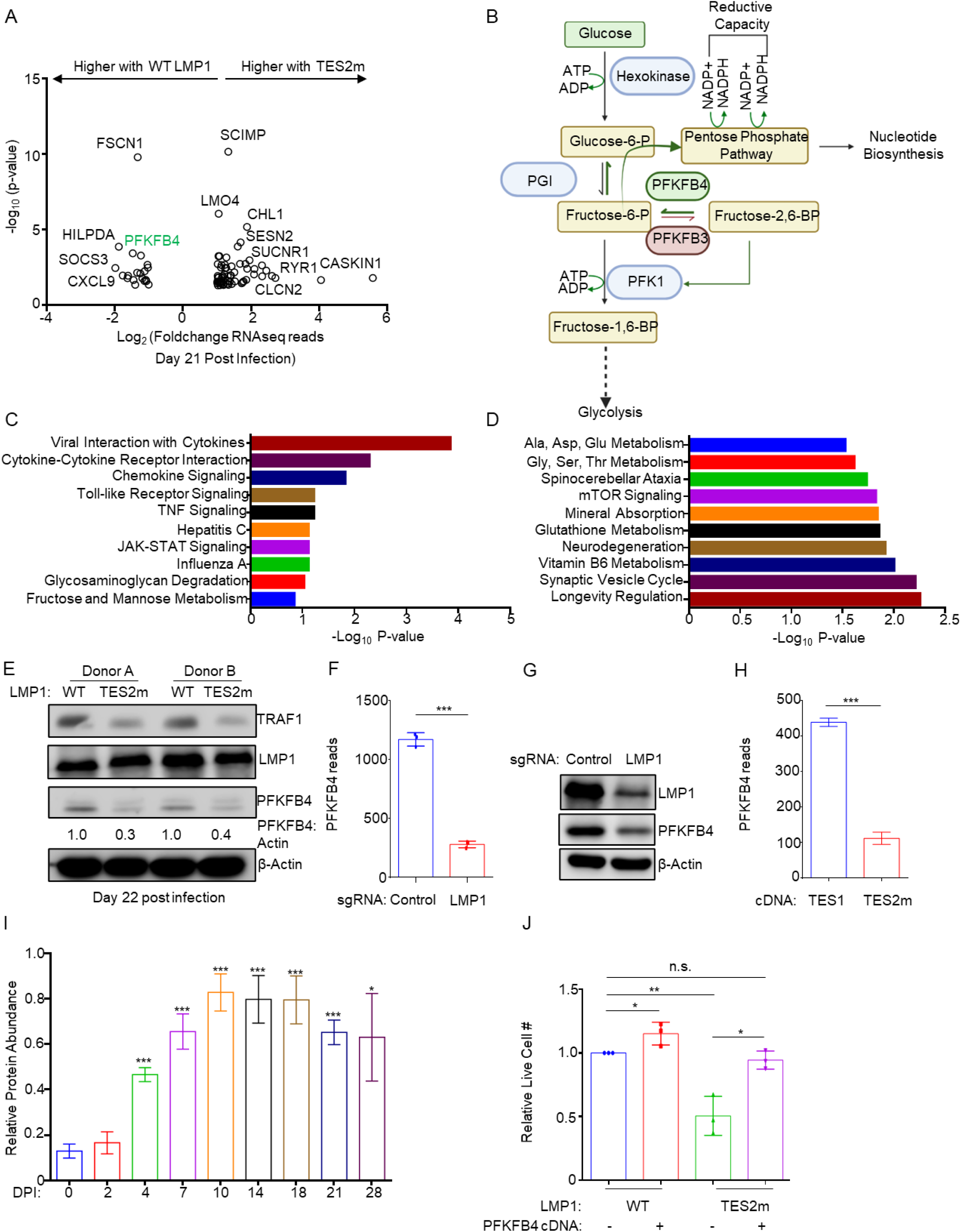
LMP1 TES2 signaling upregulates PFKFB4 to promote B cell resistance to erastin ferroptosis induction. (A) Volcano plot analysis of host transcriptome-wide genes differentially expressed in primary B cells from three independent donors infected with WT or TES2m LMP1 for 21 days. Positive fold changes indicate higher transcripts in cells with TES2m LMP1. (B) PFKFB4 glucose metabolism roles. Glucose is metabolized by glycolysis or by the PPP. Glycolysis flux is regulated via Fructose-2,6-BP, an allosteric regulator of PFK1. PFKFB3 phosphorylates Fructose-6-P into Fructose-2,6-BP while PFKFB4 phosphatase activity converts Fructose-2,6-BP to Fructose-6-P to shunt glycolytic flux to PPP. Created in BioRender. Burton, E. (2026) https://BioRender.com/i0gawc8. (C) KEGG pathway analysis of transcripts significantly upregulated in cells infected by EBV with WT LMP1 at day 21 post infection. (D) KEGG pathway analysis of transcripts significantly upregulated in cells infected by TES2m EBV at day 21 post infection. (E) PFKFB4 protein levels in WT versus TES2m LMP1 EBV infected cells. Cells from two independent donors were infected with EBV with WT LMP1 or TES2m LMP1 for 22 days. Densitometry values of PFKFB4 vs β-Actin load controls are shown. (F) Effects of LCL LMP1 KO on PFKFB4 mRNA. Median PFKFB4 reads from RNAseq analysis of Cas9+ GM12878 cells with control vs LMP1 targeting sgRNA expression for 48 hours. Shown are data from n=3 RNAseq datasets(29). (G) PFKFB4 protein levels in LMP1 KO LCLs. Cas9+ GM12878 were mock induced or induced to express LMP1 sgRNA for 48 hours. WCL was extracted and immunoblot was carried out using the indicated antibodies. (H) Effects of TES1 vs TES2 signaling inhibition on LCL PFKFB4 expression. Median PFKFB4 RNAseq reads from RNAseq analysis of Cas9+ GM12878 expressing LMP1 sgRNA together with dox-induced LMP1 TES1m vs WT LMP1 cDNA expression. Shown are data from n=3 RNAseq datasets(29). (I) Relative PFKFB4 protein abundances from proteomic analysis of primary human B-cells at the indicated days post-infection by the EBV B95.8 strain(32). Shown are mean ± SD values from n=4 replicates. (J) Effects of exogenous PFKFB4 expression on proliferation of primary B cells infected by EBV with WT versus TES2m LMP1. Peripheral blood B cells from three independent donors were infected with EBV with WT vs TES2m LMP1. After 14 days, cells were transduced with lentivirus control or with lentivirus driving stable PFKFB4 expression. Five days post-selection, cells were re-seeded. Twelve days later, live cell counts were defined. P-values were determined by one-sided Fisher’s exact test. * p<0.05, **p<0.005, ***p<0.0005. blots are representative of n=3 independent replicates.

Immunoblot analysis confirmed lower PFKFB4 expression in EBV-infected B cells with TES2m LMP1 at day 21 post-infection relative to levels in cells infected by EBV with WT LMP1. Importantly, LMP1 expression in cells infected by WT and TES2m EBV was equivalent (**Fig. 5E**). Furthermore, re-analysis of recently published RNAseq data (29) highlighted that while PFKFB4 is highly expressed in EBV-immortalized LCLs, PFKFB4 levels were strongly diminished by 24 hours post CRISPR LMP1 knockout (**Fig. 5F**). We validated that LMP1 KO in GM12878 LCLs rapidly diminished PFKFB4 levels (**Fig. 5G**). Similarly, PFKFB4 mRNAs were lower in GM12878 LCLs whose endogenous LMP1 is knocked out via CRISPR and rescued by expression of TES2m LMP1 as compared to Levels in LMP1 KO GM12878 rescued with LMP1 TES1m expression(29)(**Fig. 5H**). Notably, LMP1 and PFKFB4 levels increase with similar kinetics in newly infected peripheral blood B cells(32), reaching maximum levels by day 21 post-infection(**Fig. 5I**). This trend was confirmed via immunoblot analysis of PFKFB4 levels from whole cell lysates extracted from EBV-transforming B cells as well (**Fig. S8B**). Similar kinetics of PFKFB4 mRNA upregulation in newly infected B-cells was observed by RNAseq analyses (68)(**Fig. S8A**).

We analyzed published LCL ChIP-seq data(69–72) to gain insights into how LMP1 signaling induces PFKFB4 expression. Activating H3K4me1, H3K4me3 and H3K27ac marks were evident at the *PFKFB4* promoter (**Fig S8C**), consistent with its expression in LCLs. All five LMP1-activated NF-κB transcription factor subunits, as well as EBNA-LP, EBNA3A and to a lesser extent EBNA3C co-occupied the *PFKFB4* promoter region (**Fig S8C**).

We next asked whether PFKFB4 expression could at least partially rescue outgrowth of B cells infected by EBV with TES2m LMP1. To do so, we infected primary B cells from three independent donors infected with EBV with WT vs TES2m LMP1. At day 14 post-infection, we transduced cells with negative control lentivirus versus lentivirus stably expressing PFKFB4 cDNA, as evidenced by exogenous PFKFB4 expression (**Fig. S8D**). Successfully transduced cells were selected for five days. PFKFB4 over-expression increased growth of infected B cells with WT and TES2m LMP1 expression. Importantly, PFKFB4 expression nearly completely rescued live cell numbers of TES2m+ B-cells relative to levels in WT LMP1 infected cells (columns 1 vs 4) and significantly increased live cell numbers from levels observed at this timepoint in TES2m+ cells without PFKFB4 overexpression (**Fig. 5J**). These data are consistent with a model in which TES2 signaling drives PFKFB4 expression to support lymphoblastoid B cell immortalization at a stage where LMP1 expression reaches maximal levels.

### PFKFB4 supports LCL glutathione pools, lipid ROS levels and resistance to erastin ferroptosis induction

Although PFKFB4 is highly expressed in LCLs, its metabolism roles have not been characterized in EBV infected cells. Given our observation that PFKFB4 supports TES2-driven redox defense, we next defined the effects of PFKFB4 in two LCLs, GM12878 and GM15892. PFKFB4 depletion by either of two sgRNAs significantly increased lipid ROS levels as judged by FACS analysis of BODIPY-C11 stained cells (**Fig. 6A**). PFKFB4 depletion also significantly reduced GM12878 and GM15892 steady state GSH pools (**Fig. 6B**). PFKFB4 depletion and erastin treatment additively decreased GSH levels in both LCLs (**Fig. 6C**).

**Figure 6.**
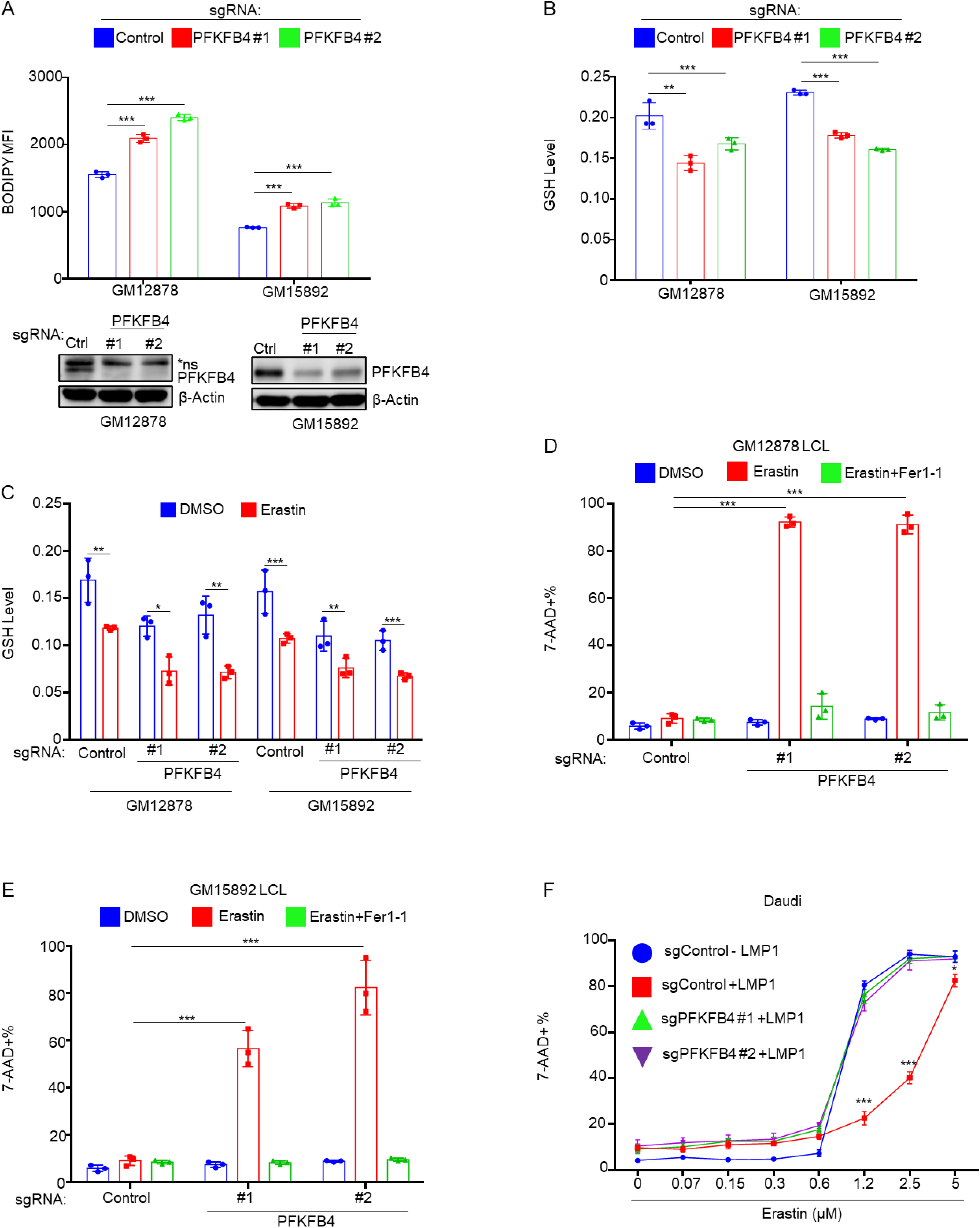
PFKFB4 protects LMP1-expressing cells from ferroptosis. (A) Lipid ROS BODIPY MFI + SD values from n=3 independent replicates of Cas9+ GM12878 or GM15892 LCLs after three days of control or PFKFB4 targeting sgRNA expression. (Bottom) Representative immunoblot of WCL from GM12878 and GM15892 LCL at this timepoint. (B) GSH levels from n=3 independent replicates of Cas9+ GM12878 and GM15892 LCLs after three days of control or PFKFB4 sgRNA expression. (C) GSH levels from n=3 independent replicates of Cas9+ GM12878 and GM15892 LCLs after two days of control or PFKFB4 sgRNA expression. Cells were then treated with DMSO or 1µM erastin for another 18 hours before GSH levels were quantitated. (D) FACS analysis of mean %7AAD+ SD values from n=3 independent replicates of Cas9+ Daudi cells after three days of control or PFKFB4 sgRNA expression. Cells were then mock induced or dox induced for LMP1 expression before treatment with indicated doses of erastin. %7-AAD+ cells were quantitated after 48 hours. (E) FACS analysis of mean %7AAD+ SD values from n=3 independent replicates of GM12878 LCL after three days of control or PFKFB4 sgRNA expression. Cells were then treated with DMSO, 1.25µM erastin and/or 2.5µM Fer-1 for 48 hours. (F) FACS analysis of mean %7AAD+ SD values from n=3 independent replicates of GM15892 LCL after three days of control or PFKFB4 sgRNA expression. Cells were then treated with DMSO, 2.5µM erastin and/or 2.5µM Fer-1 for 48 hours. P-values were determined by one-sided Fisher’s exact test. * p<0.05, **p<0.005, ***p<0.0005.

**Figure 7.**
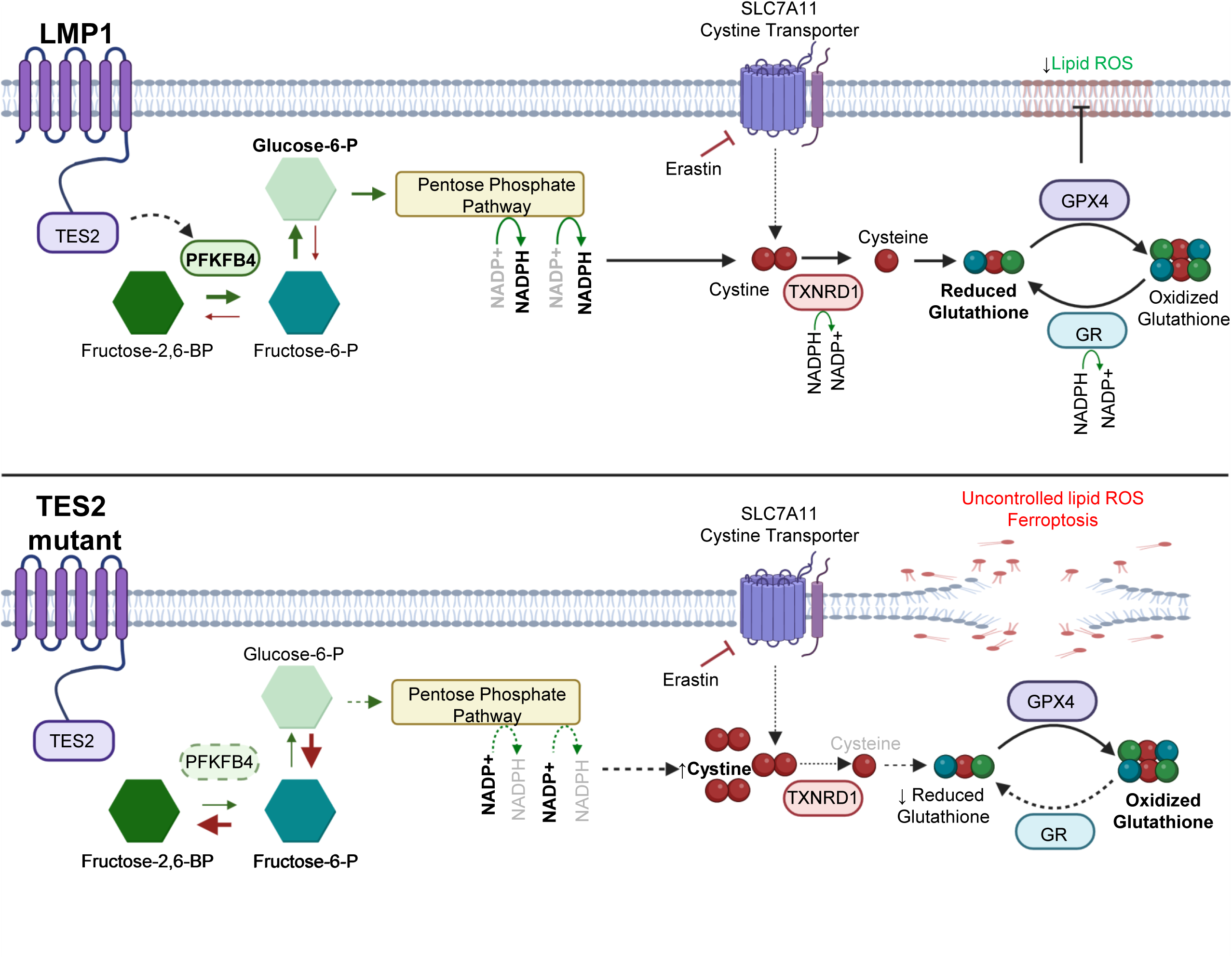
Model of LMP1 TES2-mediated, PFKFB4-dependent ferroptosis resistance. Top: LMP1 TES2 signaling promotes cystine uptake and upregulates PFKFB4, which promotes conversion of fructos-2,6 bp to Fructose-6-p. Since F-2,6-BP is a major allosteric activator of the glycolysis pathway enzyme PFK1, this serves to shunt glycolytic flux away from glycolysis and towards PPP. In turn, PPP produces NADPH to support cystine reduction to cysteine by TXNRD1 and to increase GSH pools, including by the GSSG reduction via glutathione reductase. This provides GSH for GPX4-mediated lipid ROS detoxification and ferroptosis evasion. Bottom: In the absence of TES2 signaling, EBV driven lipid metabolism produced lipid ROS can not be adequately detoxified upon blockade of cystine import by erastin due to decreased NADPH production and reduction of GSH. Consequently, erastin triggers ferroptosis induction. Created in BioRender. Burton, E. (2026) https://BioRender.com/yfk9l2n

Given the above results, we next tested whether LCLs are dependent on PFKFB4 for ferroptosis defense. We treated control or PFKFB4 depleted GM12878 or GM15892 LCLs with erastin for 24 hours. Intriguingly, PFKFB4 depletion by either sgRNA strongly sensitized LCLs to erastin-induced cell death (**Fig. 6D-E, Fig. S9A-B**). These results indicate that LCLs are highly dependent on EBV-induced PFKFB4 expression to support their cysteine and glutathione metabolism in support of redox defense. Conversely, whereas LMP1 expression protects Daudi Burkitt cells from erastin-mediated ferroptosis induction, this phenotype is dependent on PFKFB4 expression (**Fig. 6F, S9C-D**). Collectively, these results highlight PFKFB4 as a key LMP1 TES2 target that is subverted by EBV for redox defense.

## Discussion

Here, we investigated how EBV oncoprotein signaling alters redox defense to confer resistance to ferroptosis induction. Although LMP1 mimics key aspects of B cell co-receptor CD40 signaling, we found that LMP1 but not CD40 signaling engenders resistance to ferroptosis induction by erastin mediated blockade of cystine import. We provided evidence that a major LMP1 role is to support cysteine metabolism in support of reduced glutathione biosynthesis. In particular, we identified that signaling by LMP1 TES2 drives erastin resistance by upregulation of PFKFB4 levels in support of NADPH production by the pentose phosphate pathway. PFKFB4 expression was necessary for erastin resistance in Burkitt cells with conditional LMP1 expression and likewise in LCLs. Taken together with our prior observation that EBV strongly upregulates lipid metabolism and lipid ROS levels in newly infected B-cells, these results identify that a major role of LMP1 TES2 signaling is to remodel pentose phosphate pathway metabolism to boost NADPH production and cysteine metabolism in support of reduced glutathione supply for redox defense against lipid ROS.

EBV infection strongly increases lipid metabolism pathways, oxidative phosphorylation (a major source of ROS) and lipid ROS levels in newly infected human peripheral blood B cells(32, 33, 73–76). Lipid ROS reach maximum levels at day 7 post primary human B cell EBV-infection and then progressively decrease as cells transform to lymphoblastoid cell lines(41). It is noteworthy that lipid ROS levels therefore inversely correlate with LMP1 levels(32, 77). Similarly, we found that erastin-induced ferroptosis reached maximum levels at day 7 post-EBV infection and then progressively decreased, with cells refractory to erastin as they convert to LCLs. This contrasted with immunoreceptor stimulation including by CD40L and IL-4, which sensitized cells to ML-210 but not erastin driven ferroptosis induction(41). Thus, although LMP1 mimics aspects of CD40 signaling(13–15, 18, 19, 78), these data support a model in which LMP1 evolved a key role distinct is to remodel host B cell redox defense to buffer EBV oncoprotein driven lipid metabolism and lipid ROS production.

LMP1 signaling depleted cystine and increased reduced glutathione levels and drove B cell resistance to erastin SLC7A11 blockade but not to blockade of GPX4, which uses reduced glutathione as a cofactor to detoxify lipid ROS. Therefore, whereas LCLs are comparatively resistant to both SLC7A11 and GPX4 inhibition(41), TES2 is responsible for remodeling redox defense at a level upstream of GPX4, likely at the level of GSH supply. Signaling from the LMP1 C-terminal tail TES2 region was necessary and sufficient for resistance to erastin ferroptosis induction, both in Burkitt cells and in infected B cells transforming into LCLs. CD40 or TNFα signaling did not rescue Burkitt cells from erastin, nor did IKKβ inhibition, suggesting that NF-κB signaling likely does not underly this phenotype. By contrast, SMAC mimetics, which strongly induce non-canonical NF-kB did, though it also reduced Burkitt cell fitness, complicating interpretation, and CD40-induced non-canonical NF-κB failed to rescue. Furthermore, CRISPR KO of TES2 mediator TRAF6 failed to re-sensitize LMP1+ cells to erastin. Therefore, further studies are required to define the potentially novel LMP1 TES2 pathway that induces erastin resistance, presumably by upregulation of PFKFB4.

PFKFB4 expression was necessary for LMP1-induced erastin resistance. PFKFB4 is a bifunctional kinase and phosphatase that is one of four PFKFB family isoforms (PFKFB1-4, collectively referred to as phosphofructokinase 2 or PFK2). PFKFB family members have tissue-specific expression patterns and distinct kinase/phosphatase activity ratios. Of these, EBV only upregulates PFKFB4 on the protein level in newly infected B cells based on proteomic profiling(32). PFKFB4 bisphosphatase activity is relatively higher than that of other PFK2 family members(79) and PFKFB4 therefore promotes conversion of fructose-2,6-biphosphate to fructose-6 phosphate. Importantly, EBV increases glucose uptake(32, 80) and fructose-2,6-biphosphate is a major allosteric activator of the glycolysis enzyme PFK1, which drives the first committed step of the glycolysis pathway. Therefore, PFKFB4 serves to increase glycolytic flux towards PPP. It will be of interest to determine whether TES2 signaling not only induces PFKFB4 expression but also alters its phosphatase or kinase activity on the post-translational level.

Our data suggests that a major route by which LMP1 drives erastin resistance is by increasing NADPH levels. While NADPH alone does not confer redox defense, basal NADP(H) levels correlate with sensitivity to ferroptosis inducers(81).

Whereas we previously found that EBV induced one-carbon metabolism supports NADPH production in newly infected cells(32), PPP generates >60% of NADPH in most cells(82). Our metabolomic profiling suggests that LMP1 significantly increases B cell NADPH levels, likely at the level of PFKFB4. EBV also upregulates NADH/NAD levels(32), and EBNA2 induces de novo biosynthesis of nicotinamide adenine dinucleotide (NAD)(83). NADP(H) is produced through NAD(H) phosphorylation by NAD kinase(84). Therefore, EBV may have evolved multiple oncoprotein driven pathways to support NADPH levels, though PFKFB4 activity may play an obligatory role in support of reduced glutathione levels.

LMP1 upregulated NAPDH levels likely support the reduced glutathione pool by complimentary mechanisms. First, LMP1 increases expression of the cystine/glutamate antiporter SLC7A11 on the mRNA level, both when expressed alone in Burkitt cells(29) and also when co-expressed with LMP2A in germinal center B-cells in vivo(85). Consistent with this, we found that LMP1 expression increased cystine uptake. Second, LMP1 induction of PFKFB4 drives PPP to supply NADPH, used at multiple levels of redox defense. NADPH supports conversion of cystine to cysteine for incorporation into glutathione synthesis. This likely underlies the observation that LMP1 expression decreases cystine but increases glutathione levels. Furthermore, NADPH supports thioredoxin-mediated regeneration of reduced glutathione from oxidized glutathione to support GPX4-mediated redox defense. We speculate that this latter pathway is a major mechanism by which LMP1 enables erastin resistance, by recycling oxidized glutathione pools to provide reduced glutathione as the key GPX4 cofactor for lipid ROS detoxification when it cannot be synthesized de novo. It is also possible that LMP1 drives transsulfuration, in which methionine and serine metabolism produce cysteine intracellularly. It is also possible that LMP1 increases extracellular cysteine levels(30, 86), which could plausibly be imported by a neutral amino acid transporter. Alternatively, the ferroptosis suppressor FSP1/AIFM2 functions through NADPH-dependent production of ubiquinol, which is an antioxidant that can neutralize lipid ROS and that can underlie a pathway of erastin resistance(39, 87–90).

In addition to serving a key role in redox defense in B cell transformation, it is plausible that LMP1 evolved to induce PFKFB4 to boost NADPH production in particular niches in vivo, perhaps where cystine levels are limiting. For instance, rapid B cell proliferation within the germinal center microenvironment creates a high amino acid demand including for redox balance, but there is currently limited knowledge about cystine levels within secondary lymphoid microenvironments, particularly in humans. In this context, we speculate that LMP1 boosted cystine import and NADPH production helps EBV-infected B-cells growth and survival within this key microenvironment from which most B cell lymphomas arise. We note that EBV and LMP1 also activate the KEAP1/NRF2 antioxidant defense pathway(91, 92). Thus, in addition to regulation of PFKFB4, LMP1 has therefore evolved multiple pathways to alter ROS levels and redox defense.

It is noteworthy that outgrowth of primary human B cells infected by EBV with TES2 mutant LMP1 failed to grow in culture for 6 weeks unless they were grown on fibroblast feeders(25). While the authors speculated that fibroblast feeders rescued growth through cytokine signaling, a key additional role played by fibroblast feeders is to convert cystine into cysteine(31, 39, 93).

There is growing interest in targeting metabolic vulnerabilities to selectively trigger ferroptosis in cancer cells, including to overcome resistance to conventional treatments(94–97). Since LMP1 expression or the EBV latency III program(41) drive ferroptosis resistance, our studies highlight PFKFB4 as an intriguing therapeutic target. For instance, it may be possible to combine PFKFB4 inhibition with imidazole ketone erastin(98) to resensitize lymphomas with LMP1 expression to ferroptosis induction, potentially in the settings of EBV+ PTLD, diffuse large B-cell lymphoma, primary central nervous system lymphoma or Hodgkin lymphoma, all of which express LMP1. Furthermore, the PPP non-oxidative branch also strongly supports nucleotide biosynthesis through ribose-5-phosphate production, and LMP1 expression confers dependence on de novo GTP biosynthesis(62). PFKFB4 inhibition may therefore attack LMP1-expressing tumor cells by two mechanisms, starving them of NADPH for redox defense and impairing necessary nucleotide biosynthesis.

In summary, we identified that the EBV LMP1 oncogene remodels B cell metabolism pathways to support redox defense. LMP1 TES2 signaling induces PFKFB4 to confer resistance to ferroptosis induction by erastin by increasing NADPH and reduced glutathione levels. These studies identify approaches to resensitize EBV-driven lymphomas to ferroptosis.

## Acknowledgements

We sincerely thank Drs. Eric Johannsen and Makoto Ohashi for technical assistance with EBV BAC recombineering. We thank Davide Maestri for his bioinformatic analysis. Work was supported by NIH R01CA228700, R01DE033907 and P01 CA269043 to BEG. EMB and BM were supported by American Cancer Society Post-doctoral Fellowships PF-23-898493-01-TBE and PF-24-1250090-01-IBCD, respectively. The authors also acknowledge generous philanthropic support of George and Sandra K. Schussel. The funders had no role in study design, data collection and interpretation, or the decision to submit the work for publication.

## Materials and Methods

### Ethics statement

Platelet-depleted venous blood obtained from the Brigham & Women’s hospital blood bank were used for primary human B cell isolation, following our Institutional Review Board-approved protocol for discarded and de-identified samples. The Mass General Brigham Hospital Institutional Review Board (IRB) approved this study. Study approval numbers are as follows: 2022P001270 and 2018p002995. Formal consent was obtained by the Brigham & Women’s hospital blood bank during before donation.

### Cell lines and culture

293T, Daudi and Jijoye were purchased from American Type Culture Collection. P3HR-1, Daudi, and EBV-Akata [75] were obtained from Elliott Kieff., GM12878, and GM15892 LCL were obtained from Coriell. MUTU I and MUTU III were obtained from Jeff Sample and Alan Rickinson. All B-cell lines were cultured in RPMI-1640 (Invitrogen) supplemented with 10% fetal bovine serum (FBS). 293T cells were cultured in DMEM with 10% FBS. All cell lines were incubated with 1% penicillin-streptomycin (Gibco) in a humidified incubator at 37 C and 5% CO2. All cells were routinely confirmed to be mycoplasma-negative by Lonza MycoAlert assay (Lonza).

### Primary B Cell Isolation and Culture

RosetteSep and EasySep negative isolation kits (Stemcell Technologies) were used sequentially to isolate CD19+ B cells by negative selection, with the following modifications made to the manufacturer’s protocols. For RosetteSep, 40 μL antibody mixture was added per mL of blood and before Lymphoprep density medium was underlayed, prior to centrifugation. For EasySep, 10 μL antibody mixture was added per mL of B cells, followed by 15 μL magnetic bead suspension per mL of B cells. After negative selection, the cells were washed twice with 1x PBS, counted, and seeded for EBV infection studies. Cells were cultured in RPMI-1640 (Invitrogen) supplemented with 10% FBS and penicillin-streptomycin in a humidified chamber at 37 C and 5% CO2. Cells were cultured in RPMI-1640 supplemented with 10% FBS and penicillin-streptomycin in a humidified incubator at 37 C and at 5% CO2.

### BAC Recombineering

The EBV p2089 BACmid contains the complete genome of the B95.8 strain of EBV in addition to a cassette containing the prokaryotic F-factor as well as the green fluorescent protein (*GFP*) and *Hygromycin B* resistance genes in the B95.8 deletion as previously described(99). This same WT GFP-negative p2089 BACmid is the parental BACmid to all mutants in this study. LMP1 mutant BACmids were constructed using the GS1783 *E*. *coli*–based *En Passant* method previously described(100). Gene blocks were designed to mutate well-established residues responsible for LMP1 TES1 and TES2 functionality: we edited the 204PQQAT208 functional motif of the TES1 domain to AQAAA(27, 101) and mutated the 384YYD386 motif of the TES2 domain with ID(52). All BACmids were transferred to the chloramphenicol-resistant BM2710 *E*. *coli* used for infection of 293T cells for virus production. The integrity of all BACmids was confirmed by analyzing the restriction digestion patterns with multiple enzymes. All mutations were confirmed by high fidelity PCR amplification and sequencing the mutated junctions. The strategy use to generate geneblock for mutagenesis can be found in Supplemental Figure 3. A comprehensive list of primers used for confirmation of mutagenesis follows:

**Table 1.**
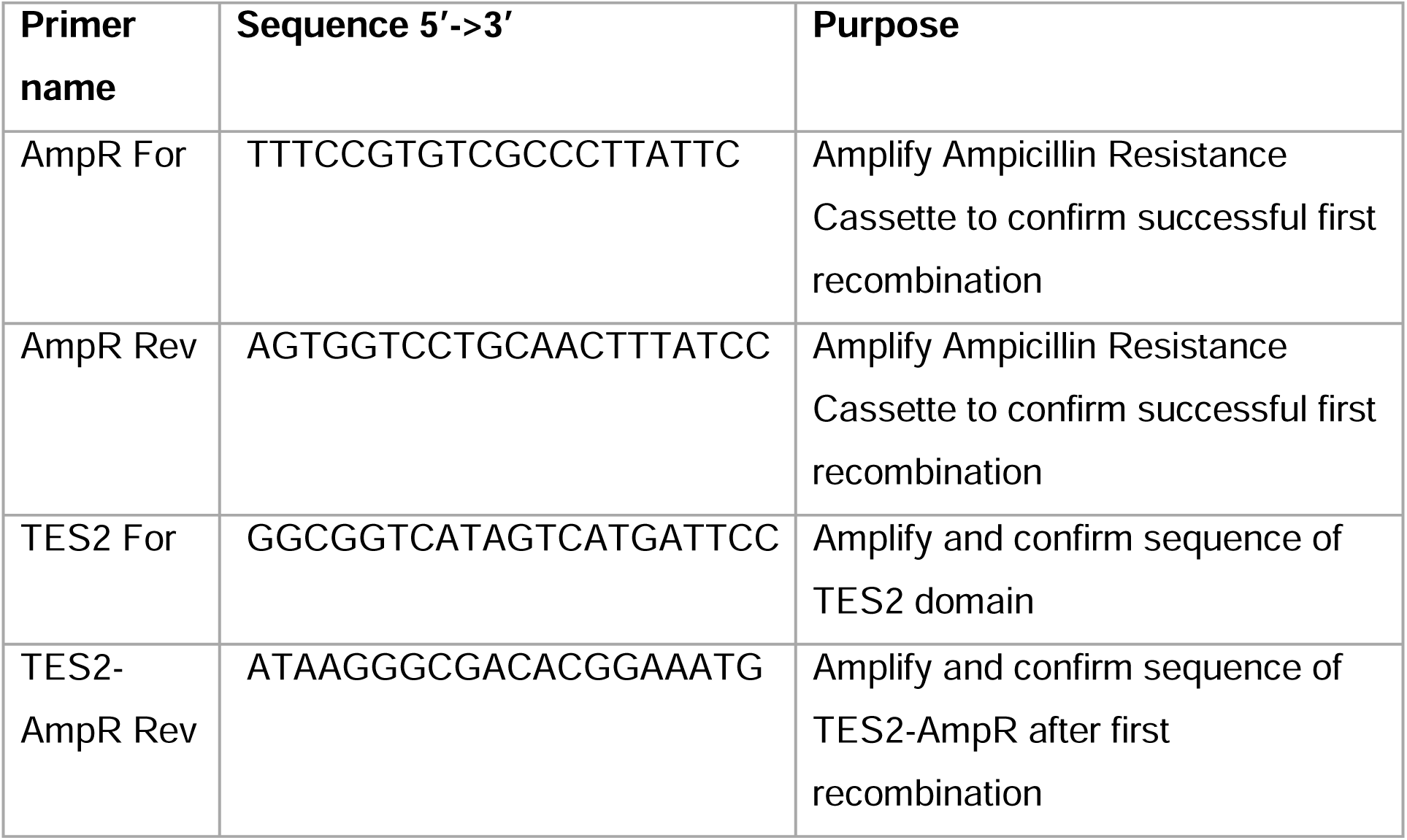
PCR primers used in BAC Recombineering recombination validation.

### Antibodies and Reagents

Antibodies against the following proteins were used in this study: β-Actin (Biolegend, 664802), PFKFB4 (Abcam, EPR28633-53), PFKFB4 (Proteintech, 29902-1-AP), EPR28633-53)XIAP (Cell Signaling Technology, # 2045S), SLC7A11/xCT (Cell Signaling Technology, 12691S) CIAP1 (Proteintech, 66626-1-IG), CIAP2 (Cell Signaling Technology, 3130s), GAPDH (EMD Millipore, MAB374), LMP1 S12 (David Thorley-Lawson), DDX1 (Bethyl, A300-521A-M), p100/p52 (EMD Millipore, 05-361), TRAF1 (Cell Signaling Biotechnology, #4715S), HA tag antibody (Cell Signaling Technology, # 3724). The following reagents were utilized in this study at the indicated concentration unless otherwise noted: DMSO (Fisher, BPBP231-100), doxycycline hyclate (Sigma, D9891-1G, 250 ng/mL), ML-210 (Cayman Chemical, 23282, as indicated), Erastin (Selleckchem, S7242, as indicated), Ferrostatin-1 (Selleckchem, S7243, 5 µM), IKK-2 inhibitor VIII (Sigma, A3485, as indicated), MEGACD40 ligand, (Enzo, ALX-522-110-C010, 50 ng/mL), Birinapant (Selleckchem, S7015, 20 µM), TNFα (Millipore, GF314, 10 ng/mL).

### Immunoblot analysis

Cells were lysed in Laemmeli buffer (0.2 M Tris-HCL, 0.4 M dithiothreitol, 277 mM SDS, 6 mM bromophenol blue, and 10% [vol/vol] glycerol) and sonicated at 4°C for 5 s using a probe sonicator at 20% amplitude and boiled at 95°C for 8 min. The whole cell lysates were resolved by 10% SDS-PAGE, transferred to nitrocellulose filters at 100 V at 4°C for 1.5 h, blocked with 5% non-fat dried milk in 1× TBST for 1 h at room temperature, and then probed with the indicated primary antibodies (diluted in 1 × TBS-T with 0.02% sodium azide) overnight at 4°C on a rotating platform. Blots were washed three times in TBST for 10 min each and then probed with horseradish peroxidase (HRP)-conjugated secondary antibodies at a dilution of 1:3,000 in 1× TBST with 5% non-fat dried milk for 1 h at room temperature. Blots were then washed three times in TBST for 10 min each, developed by ECL chemiluminescent substrate (Thermo Scientific, #34578), and imaged on Li-COR Odyssey workstation.

### B95.8 EBV Preparation and Primary B-cell Infection

B95.8 virus was prepared using 4-HT inducible ZTA system within latently-infected B95.8 cells. EBV lytic cycle was activated by treating the cells with 1 μM 4HT for 24 hours. Subsequently, the media containing 4-HT was removed and replaced with fresh RPMI1640+10%FBS+1%P/S, and the cells were cultured in RPMI 1640 medium supplemented with 10% FBS, for an additional 96 hours. The viral supernatants obtained were then cleared of producer cells by passing through a 0.45-μm filter and stored at -80 degrees C. Infectious titer of freshly thawed EBV was determined by primary B-cell transformation assay. Freshly isolated, de-identified, discarded CD19+ peripheral blood B cells were seeded in RPMI1640 with 10% FBS at a concentration of one million cells/mL for infection studies at an EBV multiplicity of infection of 0.1.

### Metabolite Extraction and Metabolomic Analysis

Metabolites were extracted according to Asara lab published protocols(102). Twenty-four hours after doxycycline (250 ng / mL, Sigma Cat#D9891-1G) addition, uninduced and LMP1-induced Burkitt cells were pelleted and resuspended in fresh RPMI/FBS. After two hours, metabolites were extracted from two million cells using 4 mL of -80 degrees C 80% methanol, made from HPLC grade water (Sigma, 270733-1L) and LC-MS-grade methanol (Fisher, A456-1). Cells were suspended in 80% methanol by vortexing and pipetting, and extraction was performed overnight in a 4 degree C room on dry ice to ensure metabolite stability. Cell debris was pelleted in a 15 mL conical at 14,000*g* for 5 minutes at 4 degrees C and supernatant was collected on dry ice. An additional round of extraction was performed using 0.5 mL of fresh 80% methanol (-80 degrees C). Cell debris were vigorously suspended using a combination of intense vortexing and pipetting with a p1000. A total of 4.5 mL of extracted metabolites were split into three 1.5 ml Eppendorf tubes and then dried using a Speedvac for ∼6-8 hours. Dried pellets were stored at -80 until resuspension in HPLC water and LC-MS/MS analysis. LC-MS/MS was performed as published(102). Peak area integrated TIC were utilized for relative comparisons of metabolites between samples.

### Targeted Mass Spectrometry

Samples were re-suspended using 20 μL HPLC grade water for mass spectrometry. 5-7 μL were injected and analyzed using a hybrid 6500 QTRAP triple quadrupole mass spectrometer (AB/SCIEX) coupled to a Prominence UFLC HPLC system (Shimadzu) via selected reaction monitoring (SRM) of a total of 298 endogenous water soluble metabolites for steady-state analyses of samples. Some metabolites were targeted in both positive and negative ion mode for a total of 309 SRM transitions using positive/negative ion polarity switching. ESI voltage was +4950V in positive ion mode and –4500V in negative ion mode. The dwell time was 3 ms per SRM transition and the total cycle time was 1.55 seconds. Approximately 9-12 data points were acquired per detected metabolite. Samples were delivered to the mass spectrometer via hydrophilic interaction chromatography (HILIC) using a 4.6 mm i.d x 10 cm Amide XBridge column (Waters) at 400 μL/min. Gradients were run starting from 85% buffer B (HPLC grade acetonitrile) to 42% B from 0-5 minutes; 42% B to 0% B from 5-16 minutes; 0% B was held from 16-24 minutes; 0% B to 85% B from 24-25 minutes; 85% B was held for 7 minutes to re-equilibrate the column. Buffer A was comprised of 20 mM ammonium hydroxide/20 mM ammonium acetate (pH=9.0) in 95:5 water:acetonitrile. Peak areas from the total ion current for each metabolite SRM transition were integrated using MultiQuant v3.0 software (AB/SCIEX).

### CRISPR/Cas9 mutagenesis

B-cell lines with stable Cas9 expression were established as described previously(103). sgRNA constructs were generated as previously described(104) using sgRNA sequences from the Broad Institute Avana or Brunello libraries. CRISPR editing was performed as previously described(105). Briefly, lentiviruses encoding sgRNAs were generated by transient transfection of 293T cells with packaging plasmids pasPAX2 (Addgene, Plasmid #12260) and pCMV-VSV-G (Addgene, Plasmid #8454). and pLentiGuide-Puro (Addgene, Plasmid #52963) plasmids. P3HR-1, Daudi, MUTU I, GM12878, and GM13111 cells stably expressing Cas9 were transduced with the lentiviruses and selected with 3 μg/mL puromycin (Thermo Fisher, Cat#A1113803) for three days before replacement with antibiotic-free media. CRISPR editing was confirmed by immunoblotting 3 days post puromycin selection. The sgRNAs used in this study were constructed using oligos based on the sequences below:

**Table 2.**
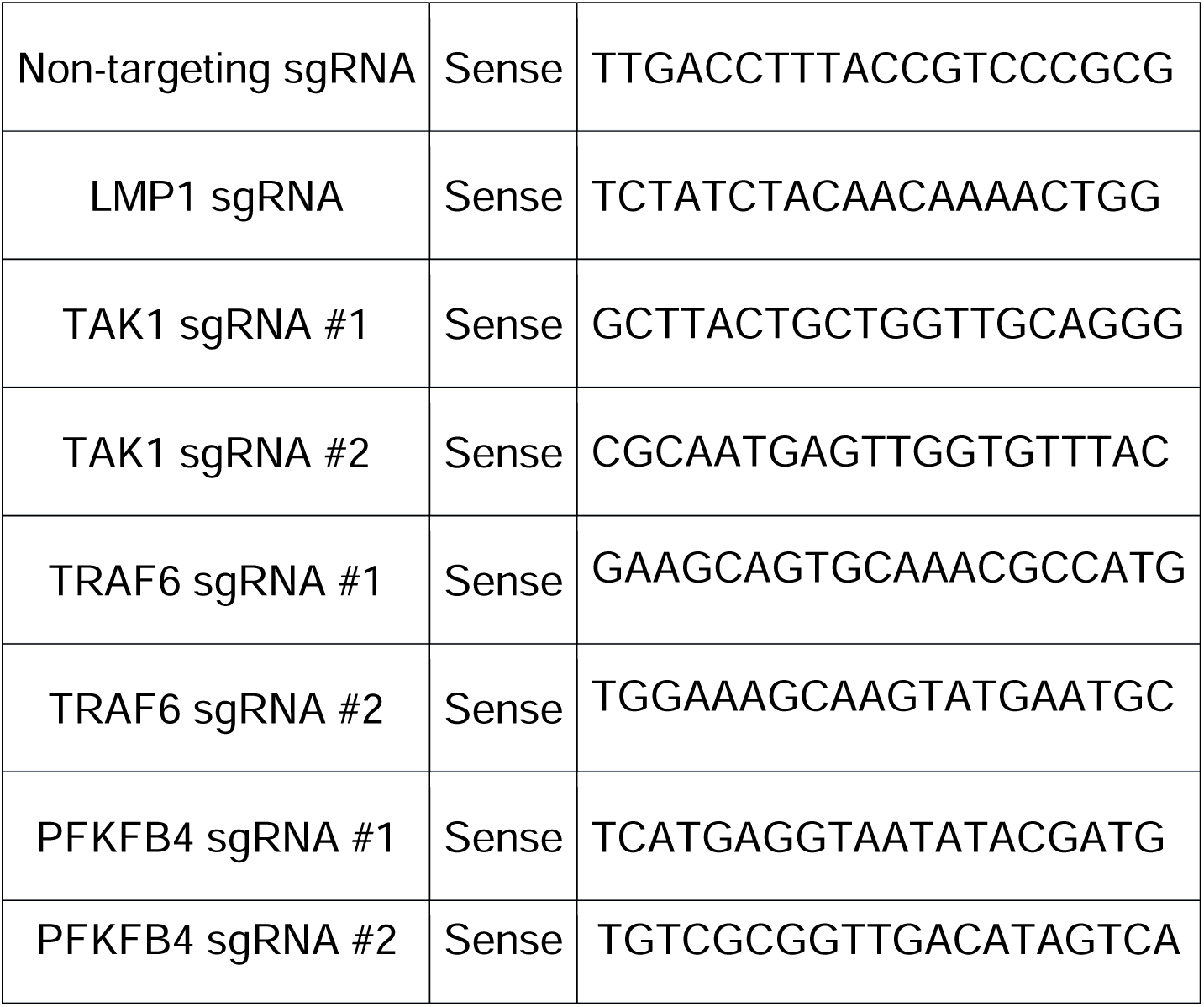
CRISPR-Cas9 sgRNA sequences.

### Flow cytometry

For 7-AAD (Thermo Fisher, Cat#A1310) viability assays, cells were harvested and washed twice with 1x PBS supplemented with 2% FBS (Gibco). Cells were then incubated with a 1 μg / mL 7-AAD solution in 1x PBS / 2% FBS for five minutes at room temperature, protected from light. Cells were then analyzed via flow cytometry. For lipid peroxidation assays, 10^5 cells were harvested, centrifuged (300 g x 5 minutes) and incubated in a u-bottom 96-well plate in 200 μL BODIPY C-11 581/591 working solution (2.5 μM BODIPY in RPMI 1640 supplemented with 10% FBS). Cells were then incubated at 37°C with 5% CO_2_ for 30 minutes, protected from light. Cells were then washed twice with 1x PBS supplemented with 2% FBS before immediate analysis via flow cytometry Labeled cells were analyzed by flow cytometry using a BD FACSCalibur instrument or Cytek Northern Lights instrument and analysis was performed with FlowJo V10.

### RNA-seq

Total RNA was isolated using RNeasy mini kit (Qiagen #74106) with in-column genomic DNA digestion step (RNase-free DNase set, Qiagen #79254) according to the manufacturer’s protocol. To construct indexed libraries, 1 µg of total RNA was used for polyA mRNA selection using NEBNext Poly(A) mRNA Magnetic Isolation Module (Cat#E7490S), and library preparation with NEBNext Ultra RNA Library Prep with Sample Purification Beads (Cat#E7765S). Each experimental treatment was performed in biological triplicate. Libraries were multi-indexed (NEB 7335L and E7500S) and pooled and sequenced on an Illumina NextSeq 500 sequencer using single 75 bp read length. Adaptor-trimmed Illumina reads for each individual library were mapped back to the human GRCh37.83 transcriptome assembly using STAR2.5.2b (<u>116</u>).

FeatureCounts was used to estimate the number of reads mapped to each contig (<u>117</u>). Only transcripts with at least five cumulative mapping counts were used in this analysis. DESeq2 was used to evaluate differential expression (DE) (<u>118</u>). DESeq2 uses a negative binomial distribution to account for overdispersion in transcriptome data sets. It is conservative and uses a heuristic approach to detect outliers while avoiding false positives. Each DE analysis was composed of a pairwise comparison between experimental group and the control group. Differentially expressed genes were identified after a correction for false discovery rate (FDR). For more stringent analyses, we set the cutoff for truly differentially expressed genes as adjusted *P* value (FDR corrected) <0.05 and absolute fold change >2. DE genes meeting this cutoff were selected and subject to downstream bioinformatics and functional analyses, including clustering, data visualization, GO annotation, and pathway analysis.

### Growth curve analysis

For growth curve analysis, cells were counted and then normalized to the same starting concentration, using the CellTiterGlo (CTG) luciferase assay (Promega, Cat#G7570). Live cell numbers were quantitated at each timepoint by CTG measurements, and values were corrected for tissue culture passage. Fold change of live cell number at each timepoint was calculated as a ratio of the value divided by the input value.

### NADP:NADPH-Glo

NADP+ and NADPH quantification was performed using NADPH-glo assay (Promega) according to the manufacturer’s protocol. 500,000 cells was used per assay, being divided up into NADP+ and NADPH extraction conditions as per protocol.

NADPH:NADP+ ratio was calculated by dividing NADPH relative luminescence units (RLUs) by NADP+ RLUs.

### Glutathione Quantification

Glutathione levels were quantified using Quantichrom Glutathione (GSH) Assay kit according to manufacturer’s specifics (BioAssay Systems, DIGT-250). One million cells were used per individual replicate.

### Cystine Uptake Assay

Cystine Uptake Assay Kit was performed according to manufacturer’s specifications. (Dojindo, up05-12).

### CellTiter-Glo

CellTiter-Glo viability assay (Promega) was performed according to the manufacturer’s protocol at the indicated time points. A total of 35 μL cells in PBS were used per assay according to manufacturer instruction.

### Software/data Presentation

Statistical analysis was assessed with Student’s t test using GraphPad Prism 7 software, where NS = not significant, p > 0.05; * p < 0.05 ** p < 0.01; *** p < 0.005. and graphs were made using GraphPad Prism 7.

---

**Fig. S1.**
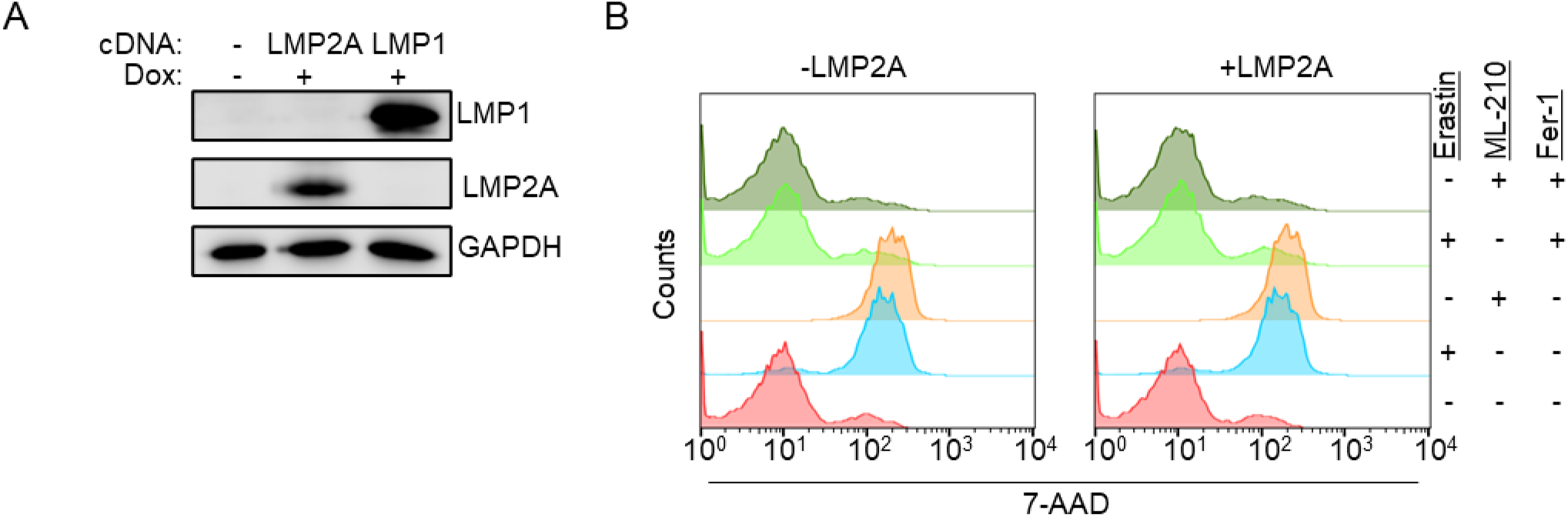

**Fig. S2.**
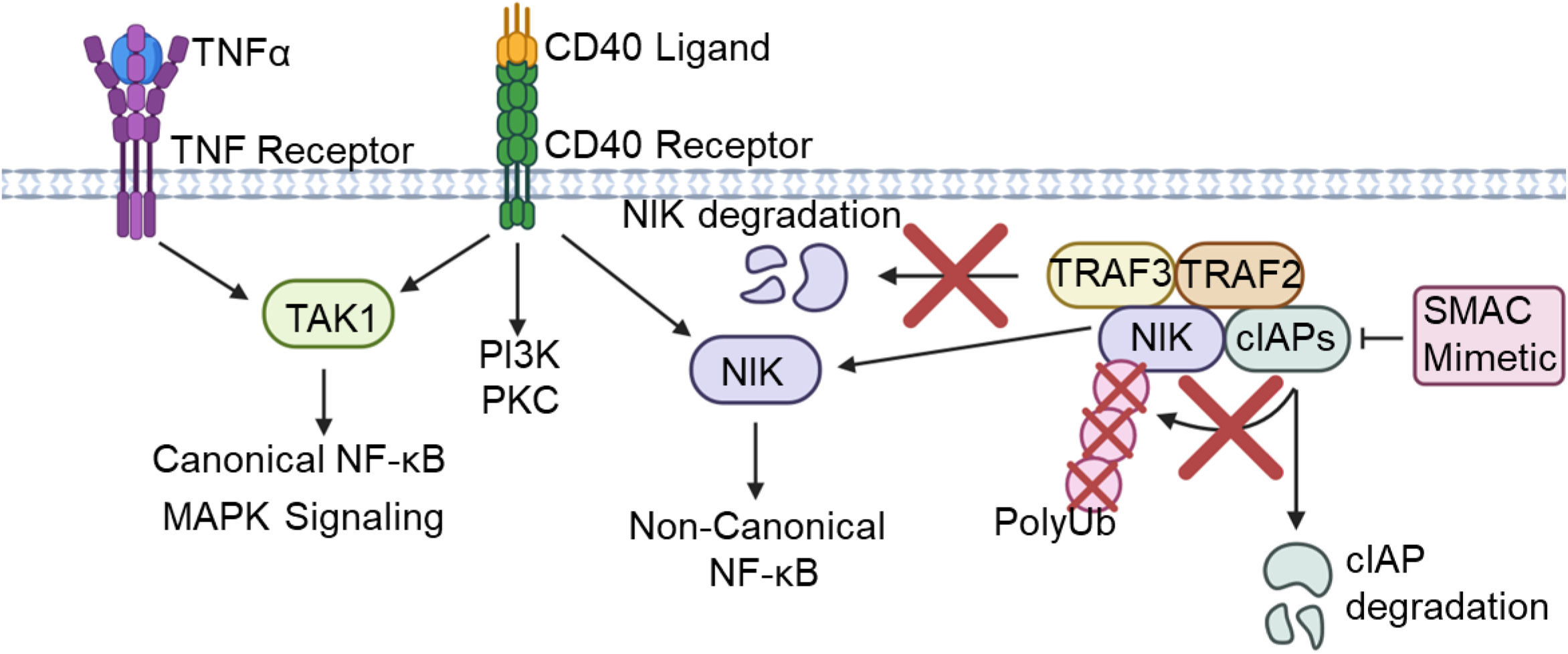

**Fig. S3.**
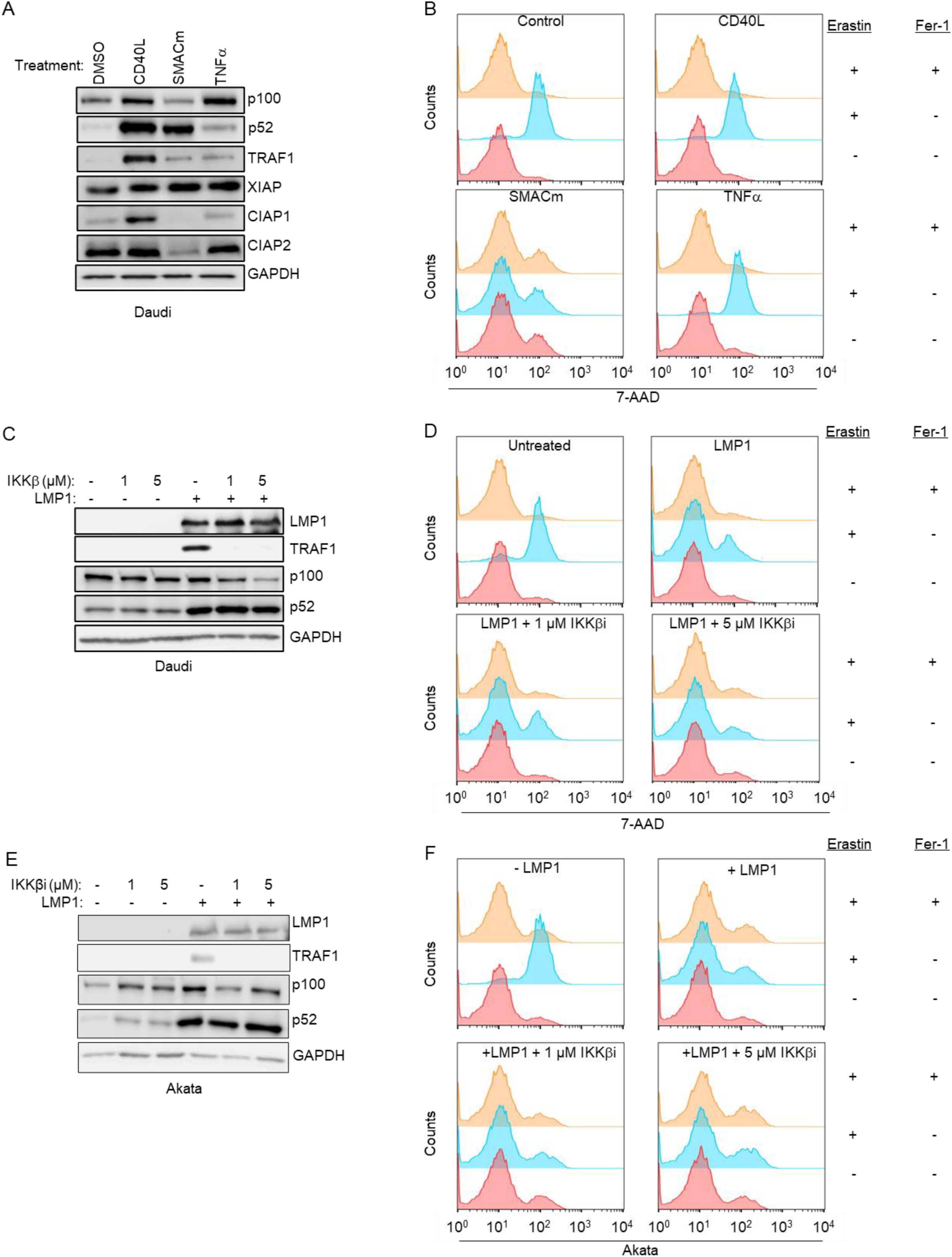

**Fig. S4.**
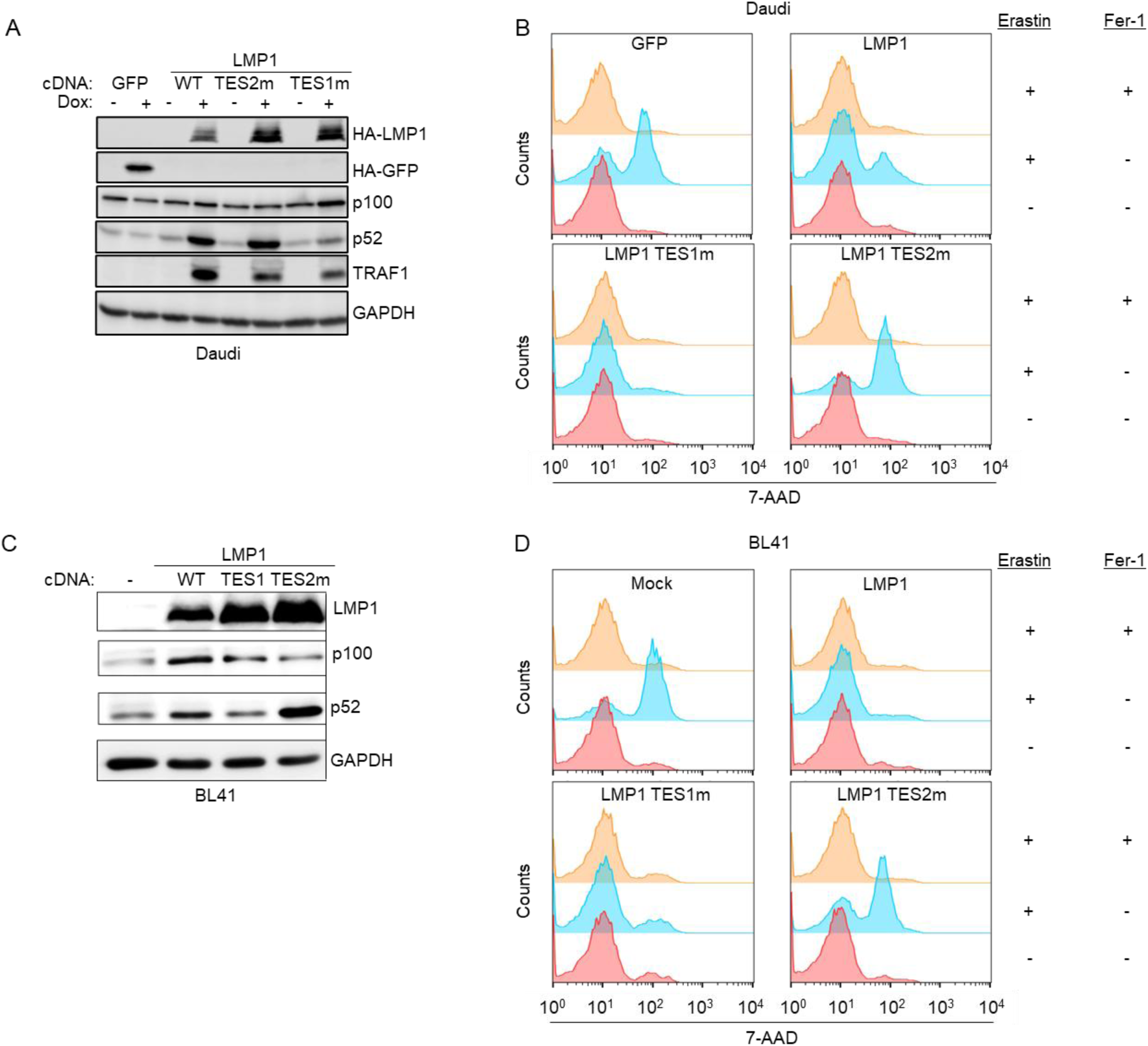

**Fig. S5.**
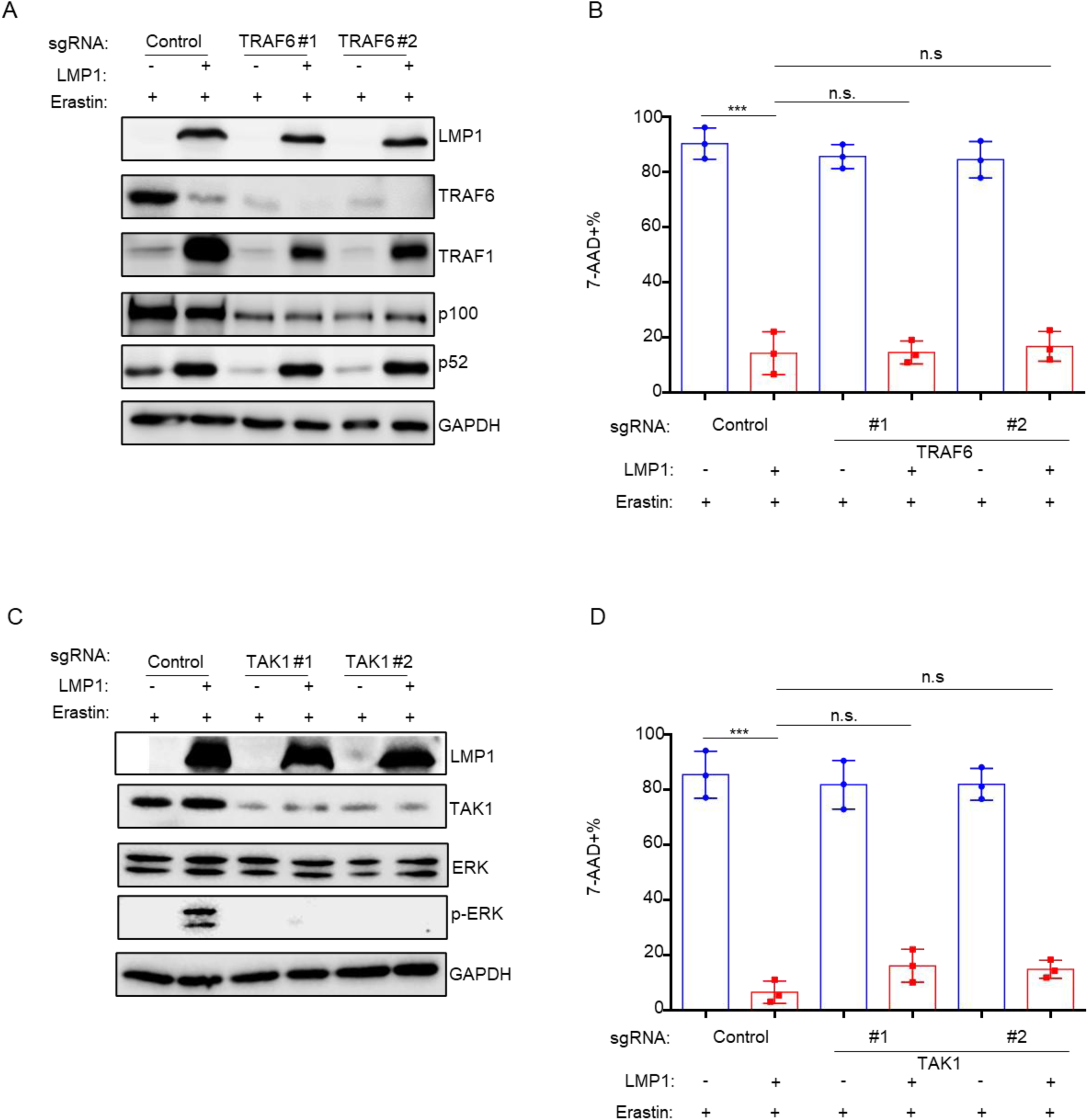

**Fig. S6.**
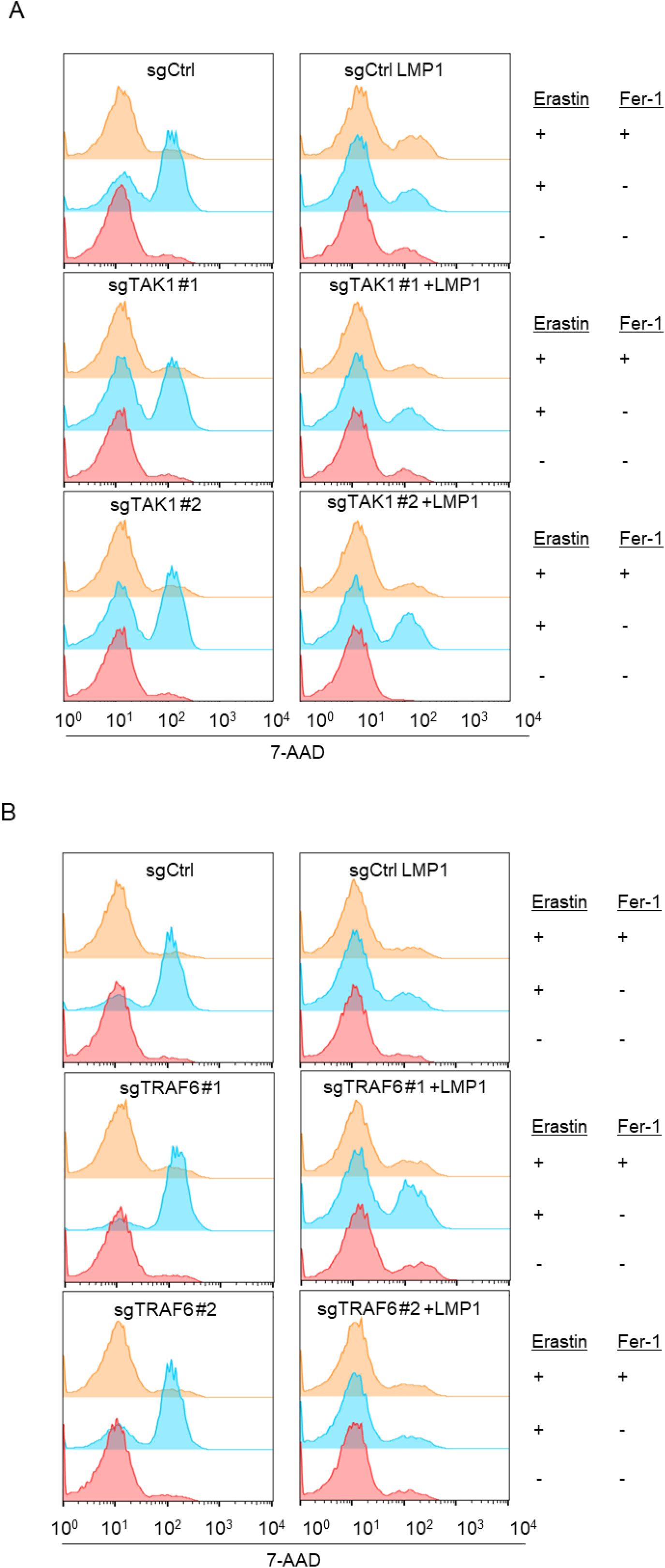

**Fig. S7.**
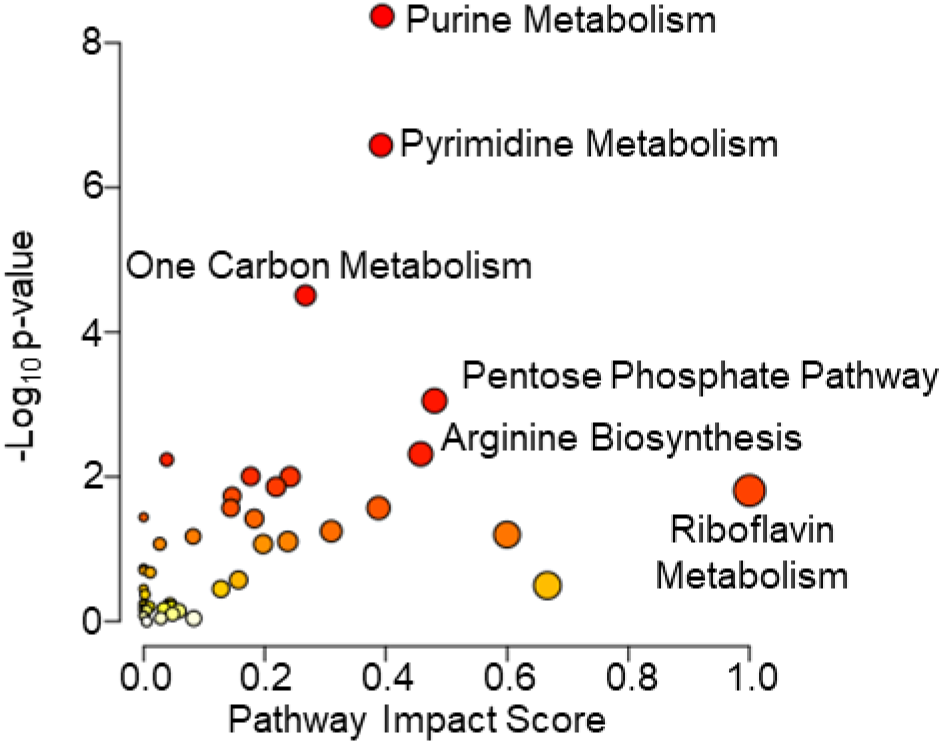

**Fig. S8.**
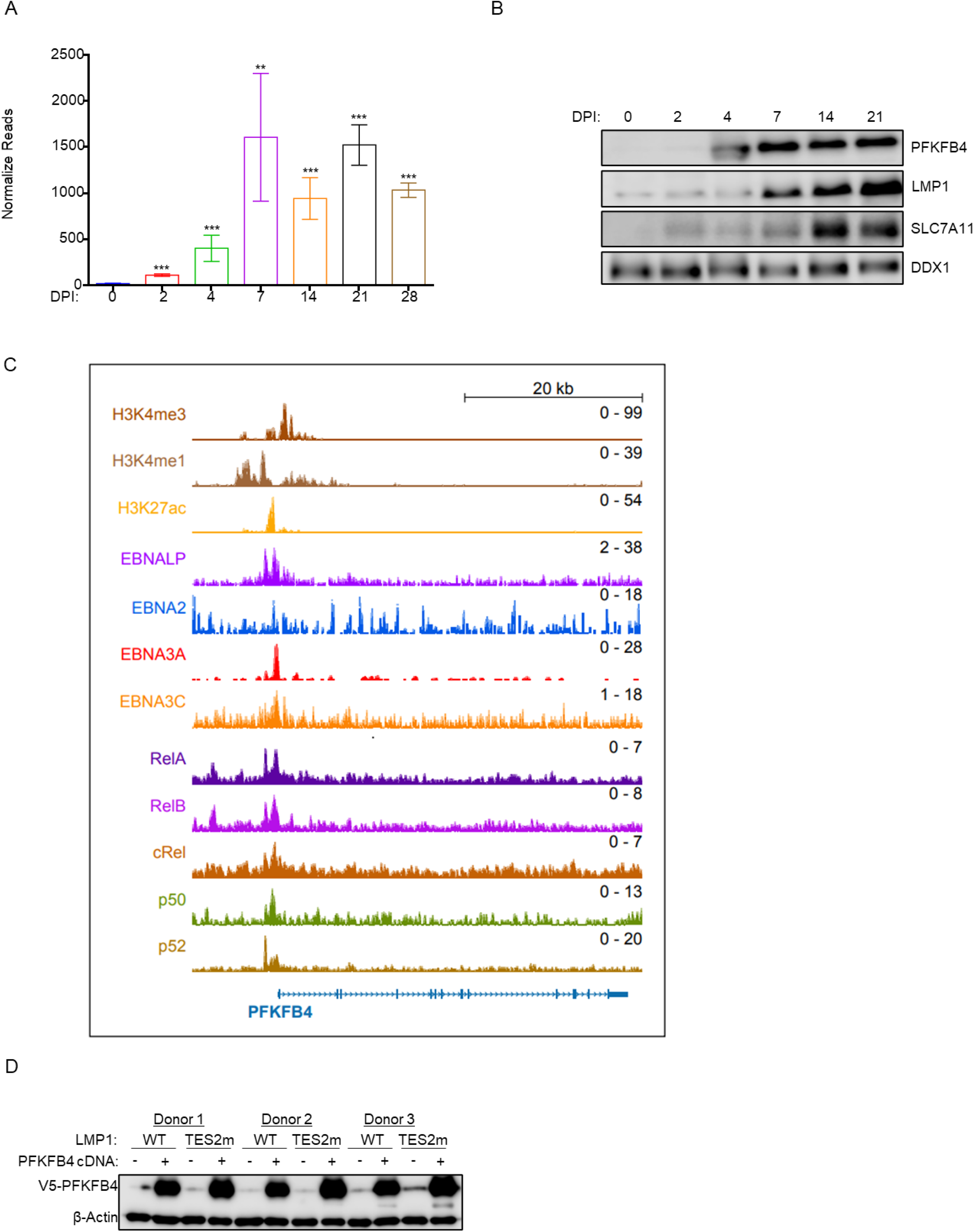

**Fig. S9.**
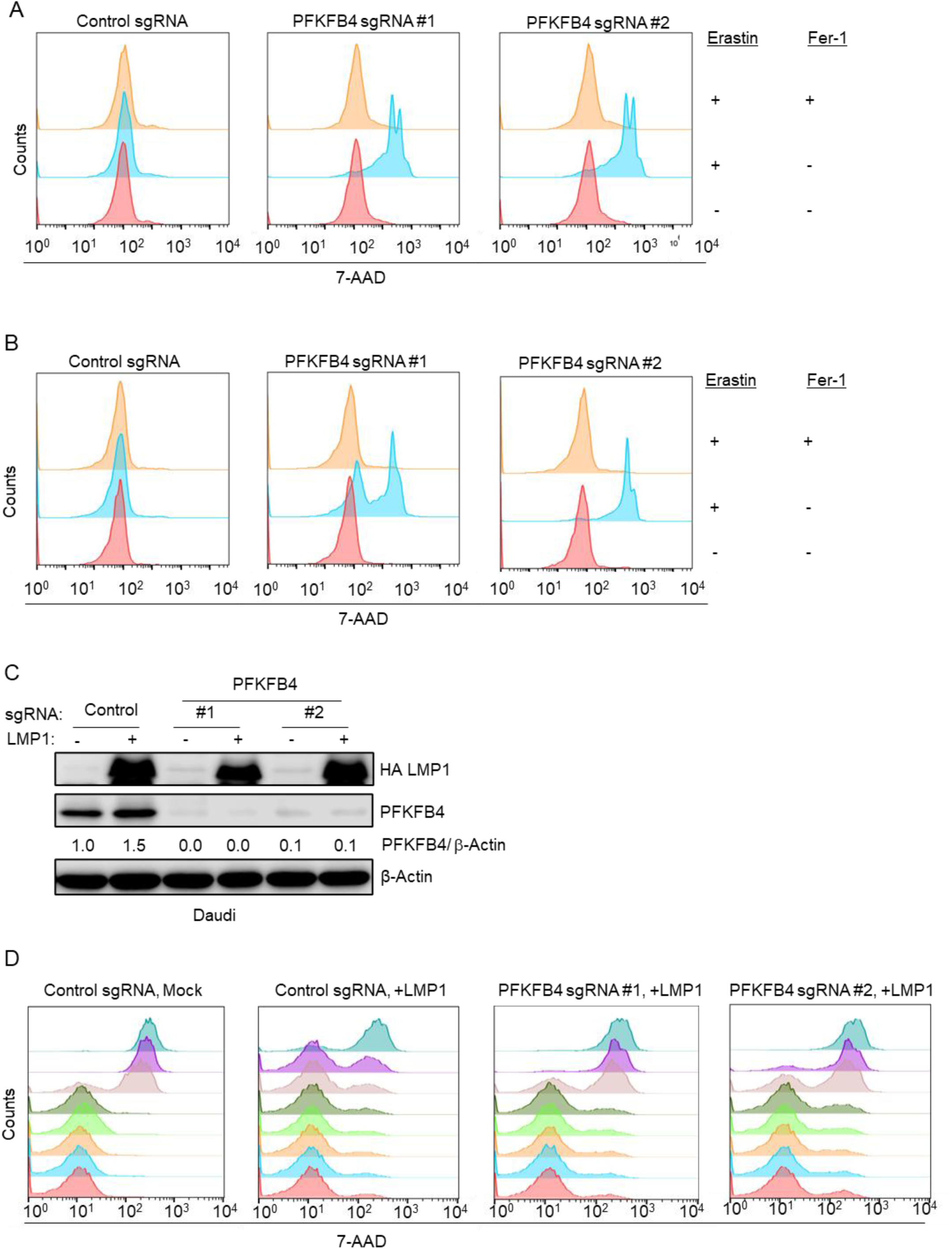

## References

1. G. Miller, The oncogenicity of Epstein-Barr virus. J Infect Dis 130, 187–205 (1974).

2. M. A. Epstein, B. G. Achong, Y. M. Barr, Virus Particles in Cultured Lymphoblasts from Burkitt’s Lymphoma. Lancet 1, 702–703 (1964).

3. Y. Pei, A. E. Lewis, E. S. Robertson, Current Progress in EBV-Associated B-Cell Lymphomas. Adv Exp Med Biol 1018, 57–74 (2017).

4. K. W. Wen, L. Wang, J. R. Menke, B. Damania, Cancers associated with human gammaherpesviruses. FEBS J 10.1111/febs.16206 (2021).

5. P. J. Farrell, Epstein-Barr Virus and Cancer. Annu Rev Pathol 14, 29–53 (2019).

6. A. Saha, E. S. Robertson, Mechanisms of B-Cell Oncogenesis Induced by Epstein-Barr Virus. J Virol 93 (2019).

7. K. Bjornevik et al., Longitudinal analysis reveals high prevalence of Epstein-Barr virus associated with multiple sclerosis. Science 375, 296–301 (2022).

8. S. S. Soldan, P. M. Lieberman, Epstein-Barr virus and multiple sclerosis. Nat Rev Microbiol 21, 51–64 (2023).

9. A. Ascherio, K. L. Munger, EBV and Autoimmunity. Curr Top Microbiol Immunol 390, 365–385 (2015).

10. S. Younis et al., Epstein-Barr virus reprograms autoreactive B cells as antigen-presenting cells in systemic lupus erythematosus. Sci Transl Med 17, eady0210 (2025).

11. M. Cortese, A. Ascherio, K. Bjornevik, EBV and Autoimmunity. Curr Top Microbiol Immunol 10.1007/82_2025_304 (2025).

12. J. E. Floettmann, M. Rowe, Epstein-Barr virus latent membrane protein-1 (LMP1) C-terminus activation region 2 (CTAR2) maps to the far C-terminus and requires oligomerisation for NF-kappaB activation. Oncogene 15, 1851–1858 (1997).

13. D. Panagopoulos, P. Victoratos, M. Alexiou, G. Kollias, G. Mosialos, Comparative analysis of signal transduction by CD40 and the Epstein-Barr virus oncoprotein LMP1 in vivo. J Virol 78, 13253–13261 (2004).

14. J. Rastelli et al., LMP1 signaling can replace CD40 signaling in B cells in vivo and has unique features of inducing class-switch recombination to IgG1. Blood 111, 1448–1455 (2008).

15. E. G. Hatzivassiliou, E. Kieff, G. Mosialos, Constitutive CD40 signaling phenocopies the transforming function of the Epstein-Barr virus oncoprotein LMP1 in vitro. Leuk Res 31, 315–320 (2007).

16. L. J. Anderson, R. Longnecker, EBV LMP2A provides a surrogate pre-B cell receptor signal through constitutive activation of the ERK/MAPK pathway. J Gen Virol 89, 1563–1568 (2008).

17. K. Fish et al., Rewiring of B cell receptor signaling by Epstein-Barr virus LMP2A. Proc Natl Acad Sci U S A 117, 26318–26327 (2020).

18. L. L. Stunz, G. A. Bishop, Latent membrane protein 1 and the B lymphocyte-a complex relationship. Crit Rev Immunol 34, 177–198 (2014).

19. L. W. Wang, S. Jiang, B. E. Gewurz, Epstein-Barr Virus LMP1-Mediated Oncogenicity. J Virol 91 (2017).

20. A. Kieser, K. R. Sterz, The Latent Membrane Protein 1 (LMP1). Curr Top Microbiol Immunol 391, 119–149 (2015).

21. L. Wang, S. Ning, New Look of EBV LMP1 Signaling Landscape. Cancers (Basel) 13 (2021).

22. B. A. Mainou, D. N. Everly, Jr., N. Raab-Traub, Unique signaling properties of CTAR1 in LMP1-mediated transformation. J Virol 81, 9680–9692 (2007).

23. D. S. Huen, S. A. Henderson, D. Croom-Carter, M. Rowe, The Epstein-Barr virus latent membrane protein-1 (LMP1) mediates activation of NF-kappa B and cell surface phenotype via two effector regions in its carboxy-terminal cytoplasmic domain. Oncogene 10, 549–560 (1995).

24. K. M. Kaye, K. M. Izumi, E. Kieff, Epstein-Barr virus latent membrane protein 1 is essential for B-lymphocyte growth transformation. Proc Natl Acad Sci U S A 90, 9150–9154 (1993).

25. K. M. Kaye, K. M. Izumi, G. Mosialos, E. Kieff, The Epstein-Barr virus LMP1 cytoplasmic carboxy terminus is essential for B-lymphocyte transformation; fibroblast cocultivation complements a critical function within the terminal 155 residues. J Virol 69, 675–683 (1995).

26. O. Devergne et al., Association of TRAF1, TRAF2, and TRAF3 with an Epstein-Barr virus LMP1 domain important for B-lymphocyte transformation: role in NF-kappaB activation. Mol Cell Biol 16, 7098–7108 (1996).

27. K. M. Izumi, K. M. Kaye, E. D. Kieff, The Epstein-Barr virus LMP1 amino acid sequence that engages tumor necrosis factor receptor associated factors is critical for primary B lymphocyte growth transformation. Proc Natl Acad Sci U S A 94, 1447–1452 (1997).

28. K. M. Kaye et al., An Epstein-Barr virus that expresses only the first 231 LMP1 amino acids efficiently initiates primary B-lymphocyte growth transformation. J Virol 73, 10525–10530 (1999).

29. B. Mitra et al., Characterization of target gene regulation by the two Epstein-Barr virus oncogene LMP1 domains essential for B-cell transformation. mBio 14, e0233823 (2023).

30. A. Banjac et al., The cystine/cysteine cycle: a redox cycle regulating susceptibility versus resistance to cell death. Oncogene 27, 1618–1628 (2008).

31. S. Bannai, T. Ishii, Formation of sulfhydryl groups in the culture medium by human diploid fibroblasts. J Cell Physiol 104, 215–223 (1980).

32. L. W. Wang et al., Epstein-Barr-Virus-Induced One-Carbon Metabolism Drives B Cell Transformation. Cell Metab 30, 539–555 e511 (2019).

33. L. W. Wang et al., Epstein-Barr virus subverts mevalonate and fatty acid pathways to promote infected B-cell proliferation and survival. PLoS Pathog 15, e1008030 (2019).

34. S. J. Dixon et al., Ferroptosis: an iron-dependent form of nonapoptotic cell death. Cell 149, 1060–1072 (2012).

35. B. R. Stockwell et al., Ferroptosis: A Regulated Cell Death Nexus Linking Metabolism, Redox Biology, and Disease. Cell 171, 273–285 (2017).

36. W. S. Yang et al., Regulation of ferroptotic cancer cell death by GPX4. Cell 156, 317–331 (2014).

37. C. D. Poltorack, S. J. Dixon, Understanding the role of cysteine in ferroptosis: progress & paradoxes. FEBS J 10.1111/febs.15842 (2021).

38. E. M. Burton, J. Voyer, B. E. Gewurz, Epstein-Barr virus latency programs dynamically sensitize B-cells to ferroptosis. PNAS, Submitted 10/01/2021 (2021).

39. G. W. Bornkamm, G. L. Kelly, A. M. Ross, Burkitt’s Lymphoma and Early B Cell Transformation as Paradigms of How Epstein-Barr Virus Overcomes Apoptosis and Ferroptosis. Curr Top Microbiol Immunol 10.1007/82_2025_301 (2025).

40. M. Brielmeier et al., Cloning of phospholipid hydroperoxide glutathione peroxidase (PHGPx) as an anti-apoptotic and growth promoting gene of Burkitt lymphoma cells. Biofactors 14, 179–190 (2001).

41. E. M. Burton, J. Voyer, B. E. Gewurz, Epstein-Barr virus latency programs dynamically sensitize B cells to ferroptosis. Proc Natl Acad Sci U S A 119, e2118300119 (2022).

42. M. Weiwer et al., Development of small-molecule probes that selectively kill cells induced to express mutant RAS. Bioorg Med Chem Lett 22, 1822–1826 (2012).

43. S. Dolma, S. L. Lessnick, W. C. Hahn, B. R. Stockwell, Identification of genotype-selective antitumor agents using synthetic lethal chemical screening in engineered human tumor cells. Cancer Cell 3, 285–296 (2003).

44. G. Miotto et al., Insight into the mechanism of ferroptosis inhibition by ferrostatin-1. Redox Biol 28, 101328 (2020).

45. E. D. Cahir-McFarland et al., Role of NF-kappa B in cell survival and transcription of latent membrane protein 1-expressing or Epstein-Barr virus latency III-infected cells. J Virol 78, 4108–4119 (2004).

46. B. E. Gewurz et al., Canonical NF-kappaB activation is essential for Epstein-Barr virus latent membrane protein 1 TES2/CTAR2 gene regulation. J Virol 85, 6764–6773 (2011).

47. Y. J. Song, K. M. Izumi, N. P. Shinners, B. E. Gewurz, E. Kieff, IRF7 activation by Epstein-Barr virus latent membrane protein 1 requires localization at activation sites and TRAF6, but not TRAF2 or TRAF3. Proc Natl Acad Sci U S A 105, 18448–18453 (2008).

48. B. A. Mainou, D. N. Everly, Jr., N. Raab-Traub, Epstein-Barr virus latent membrane protein 1 CTAR1 mediates rodent and human fibroblast transformation through activation of PI3K. Oncogene 24, 6917–6924 (2005).

49. C. P. Kung, D. G. Meckes, Jr., N. Raab-Traub, Epstein-Barr virus LMP1 activates EGFR, STAT3, and ERK through effects on PKCdelta. J Virol 85, 4399–4408 (2011).

50. A. Kieser et al., Epstein-Barr virus latent membrane protein-1 triggers AP-1 activity via the c-Jun N-terminal kinase cascade. EMBO J 16, 6478–6485 (1997).

51. C. Krepler et al., The novel SMAC mimetic birinapant exhibits potent activity against human melanoma cells. Clin Cancer Res 19, 1784–1794 (2013).

52. K. M. Izumi, E. D. Kieff, The Epstein-Barr virus oncogene product latent membrane protein 1 engages the tumor necrosis factor receptor-associated death domain protein to mediate B lymphocyte growth transformation and activate NF-kappaB. Proc Natl Acad Sci U S A 94, 12592–12597 (1997).

53. K. M. Izumi et al., The residues between the two transformation effector sites of Epstein-Barr virus latent membrane protein 1 are not critical for B-lymphocyte growth transformation. J Virol 73, 9908–9916 (1999).

54. K. M. Izumi et al., The Epstein-Barr virus oncoprotein latent membrane protein 1 engages the tumor necrosis factor receptor-associated proteins TRADD and receptor-interacting protein (RIP) but does not induce apoptosis or require RIP for NF-kappaB activation. Mol Cell Biol 19, 5759–5767 (1999).

55. F. Schneider et al., The viral oncoprotein LMP1 exploits TRADD for signaling by masking its apoptotic activity. PLoS Biol 6, e8 (2008).

56. A. Kieser, C. Kaiser, W. Hammerschmidt, LMP1 signal transduction differs substantially from TNF receptor 1 signaling in the molecular functions of TRADD and TRAF2. EMBO J 18, 2511–2521 (1999).

57. A. Shkoda et al., The germinal center kinase TNIK is required for canonical NF-kappaB and JNK signaling in B-cells by the EBV oncoprotein LMP1 and the CD40 receptor. PLoS Biol 10, e1001376 (2012).

58. L. E. Huye, S. Ning, M. Kelliher, J. S. Pagano, Interferon regulatory factor 7 is activated by a viral oncoprotein through RIP-dependent ubiquitination. Mol Cell Biol 27, 2910–2918 (2007).

59. S. Ning, J. S. Pagano, The A20 deubiquitinase activity negatively regulates LMP1 activation of IRF7. J Virol 84, 6130–6138 (2010).

60. K. M. Kaye et al., Tumor necrosis factor receptor associated factor 2 is a mediator of NF-kappa B activation by latent infection membrane protein 1, the Epstein-Barr virus transforming protein. Proc Natl Acad Sci U S A 93, 11085–11090 (1996).

61. C. D. Gregory, M. Rowe, A. B. Rickinson, Different Epstein-Barr virus-B cell interactions in phenotypically distinct clones of a Burkitt’s lymphoma cell line. J Gen Virol 71 **(Pt** **7**), 1481–1495 (1990).

62. E. M. Burton et al., Epstein-Barr virus latent membrane protein 1 subverts IMPDH pathways to drive B-cell oncometabolism. PLoS Pathog 21, e1013092 (2025).

63. I. Pader et al., Thioredoxin-related protein of 14 kDa is an efficient L-cystine reductase and S-denitrosylase. Proc Natl Acad Sci U S A 111, 6964–6969 (2014).

64. C. Pan et al., Sestrin2 as a gatekeeper of cellular homeostasis: Physiological effects for the regulation of hypoxia-related diseases. J Cell Mol Med 25, 5341–5350 (2021).

65. B. Y. Shin, S. H. Jin, I. J. Cho, S. H. Ki, Nrf2-ARE pathway regulates induction of Sestrin-2 expression. Free Radic Biol Med 53, 834–841 (2012).

66. S. H. Ro et al., Sestrin2 inhibits uncoupling protein 1 expression through suppressing reactive oxygen species. Proc Natl Acad Sci U S A 111, 7849–7854 (2014).

67. S. Ros et al., Functional metabolic screen identifies 6-phosphofructo-2-kinase/fructose-2,6-biphosphatase 4 as an important regulator of prostate cancer cell survival. Cancer Discov 2, 328–343 (2012).

68. C. Wang et al., RNA Sequencing Analyses of Gene Expression during Epstein-Barr Virus Infection of Primary B Lymphocytes. J Virol 93 (2019).

69. E. P. Consortium, The ENCODE (ENCyclopedia Of DNA Elements) Project. Science 306, 636–640 (2004).

70. B. Zhao et al., The NF-kappaB genomic landscape in lymphoblastoid B cells. Cell Rep 8, 1595–1606 (2014).

71. B. Zhao et al., Epstein-Barr virus exploits intrinsic B-lymphocyte transcription programs to achieve immortal cell growth. Proc Natl Acad Sci U S A 108, 14902–14907 (2011).

72. S. C. Schmidt et al., Epstein-Barr virus nuclear antigen 3A partially coincides with EBNA3C genome-wide and is tethered to DNA through BATF complexes. Proc Natl Acad Sci U S A 112, 554–559 (2015).

73. L. Havey, H. You, H. Xian, J. M. Asara, R. Guo, Epstein-Barr virus-transformed B-cells from a hypoxia model of the germinal center requires external unsaturated fatty acids. PLoS Pathog 21, e1013694 (2025).

74. E. N. Bonglack et al., Fatty acid desaturases link cell metabolism pathways to promote proliferation of Epstein-Barr virus-infected B cells. PLoS Pathog 21, e1012685 (2025).

75. K. L. Magon, J. L. Parish, From infection to cancer: how DNA tumour viruses alter host cell central carbon and lipid metabolism. Open Biol 11, 210004 (2021).

76. M. Hulse, S. M. Johnson, S. Boyle, L. B. Caruso, I. Tempera, Epstein-Barr Virus-Encoded Latent Membrane Protein 1 and B-Cell Growth Transformation Induce Lipogenesis through Fatty Acid Synthase. J Virol 95 (2021).

77. A. M. Price, J. E. Messinger, M. A. Luftig, c-Myc Represses Transcription of Epstein-Barr Virus Latent Membrane Protein 1 Early after Primary B Cell Infection. J Virol 92 (2018).

78. A. Kieser, The Latent Membrane Protein 1 (LMP1): Biological Functions and Molecular Mechanisms. Curr Top Microbiol Immunol 10.1007/82_2025_321 (2025).

79. M. Yi et al., 6-Phosphofructo-2-kinase/fructose-2,6-biphosphatase 3 and 4: A pair of valves for fine-tuning of glucose metabolism in human cancer. Mol Metab 20, 1–13 (2019).

80. K. McFadden et al., Metabolic stress is a barrier to Epstein-Barr virus-mediated B-cell immortalization. Proc Natl Acad Sci U S A 113, E782–790 (2016).

81. K. Shimada, M. Hayano, N. C. Pagano, B. R. Stockwell, Cell-Line Selectivity Improves the Predictive Power of Pharmacogenomic Analyses and Helps Identify NADPH as Biomarker for Ferroptosis Sensitivity. Cell Chem Biol 23, 225–235 (2016).

82. G. Kroemer, J. Pouyssegur, Tumor cell metabolism: cancer’s Achilles’ heel. Cancer Cell 13, 472–482 (2008).

83. B. Muller-Durovic et al., A metabolic dependency of EBV can be targeted to hinder B cell transformation. Science 385, eadk4898 (2024).

84. N. Pollak, M. Niere, M. Ziegler, NAD kinase levels control the NADPH concentration in human cells. J Biol Chem 282, 33562–33571 (2007).

85. T. Minamitani et al., Mouse model of Epstein-Barr virus LMP1- and LMP2A-driven germinal center B-cell lymphoproliferative disease. Proc Natl Acad Sci U S A 114, 4751–4756 (2017).

86. P. K. Mandal et al., System x(c)- and thioredoxin reductase 1 cooperatively rescue glutathione deficiency. J Biol Chem 285, 22244–22253 (2010).

87. C. Mao et al., DHODH-mediated ferroptosis defence is a targetable vulnerability in cancer. Nature 593, 586–590 (2021).

88. P. Koppula et al., A targetable CoQ-FSP1 axis drives ferroptosis- and radiation-resistance in KEAP1 inactive lung cancers. Nat Commun 13, 2206 (2022).

89. S. Zhang et al., FSP1 oxidizes NADPH to suppress ferroptosis. Cell Res 33, 967–970 (2023).

90. B. B. Silver, C. Murphy, E. J. Tokar, B. K. Sinha, Ferroptosis Suppressor Protein 1 (FSP1)-CoQ10-NADPH-Axis Is Responsible for Erastin Resistance in MCF-7 Breast Cancer Cells. Antioxidants (Basel) 15 (2026).

91. L. Wang et al., p62-mediated Selective autophagy endows virus-transformed cells with insusceptibility to DNA damage under oxidative stress. PLoS Pathog 15, e1007541 (2019).

92. L. Wang et al., The master antioxidant defense is activated during EBV latent infection. J Virol 97, e0095323 (2023).

93. T. Ishii, I. Hishinuma, S. Bannai, Y. Sugita, Mechanism of growth promotion of mouse lymphoma L1210 cells in vitro by feeder layer or 2-mercaptoethanol. J Cell Physiol 107, 283–293 (1981).

94. J. M. Ubellacker, S. J. Dixon, Prospects for ferroptosis therapies in cancer. Nat Cancer 6, 1326–1336 (2025).

95. A. Wahida, M. Conrad, Decoding ferroptosis for cancer therapy. Nat Rev Cancer 25, 910–924 (2025).

96. A. R. Brown, T. Hirschhorn, B. R. Stockwell, Ferroptosis-disease perils and therapeutic promise. Science 386, 848–849 (2024).

97. C. Mao, D. Jiang, A. C. Koong, B. Gan, Exploiting metabolic cell death for cancer therapy. Nat Rev Cancer 26, 27–45 (2026).

98. Y. Zhang et al., Imidazole Ketone Erastin Induces Ferroptosis and Slows Tumor Growth in a Mouse Lymphoma Model. Cell Chem Biol 26, 623–633 e629 (2019).

99. H. J. Delecluse, T. Hilsendegen, D. Pich, R. Zeidler, W. Hammerschmidt, Propagation and recovery of intact, infectious Epstein-Barr virus from prokaryotic to human cells. Proc Natl Acad Sci U S A 95, 8245–8250 (1998).

100. B. K. Tischer, G. A. Smith, N. Osterrieder, En passant mutagenesis: a two step markerless red recombination system. Methods Mol Biol 634, 421–430 (2010).

101. O. Devergne et al., Role of the TRAF binding site and NF-kappaB activation in Epstein-Barr virus latent membrane protein 1-induced cell gene expression. J Virol 72, 7900–7908 (1998).

102. M. Yuan, S. B. Breitkopf, X. Yang, J. M. Asara, A positive/negative ion-switching, targeted mass spectrometry-based metabolomics platform for bodily fluids, cells, and fresh and fixed tissue. Nat Protoc 7, 872–881 (2012).

103. S. Jiang et al., CRISPR/Cas9-Mediated Genome Editing in Epstein-Barr Virus-Transformed Lymphoblastoid B-Cell Lines. Curr Protoc Mol Biol 121, 31 12 31–31 12 23 (2018).

104. J. G. Doench et al., Optimized sgRNA design to maximize activity and minimize off-target effects of CRISPR-Cas9. Nat Biotechnol 34, 184–191 (2016).

105. Y. Ma et al., CRISPR/Cas9 Screens Reveal Epstein-Barr Virus-Transformed B Cell Host Dependency Factors. Cell Host Microbe 21, 580–591 e587 (2017).

