## Supplemental Figure Legends for "Epstein-Barr Virus Latent Membrane Protein 1 Suppresses Ferroptosis via Pentose Phosphate Pathway and Glutathione Metabolism"

**Figure S1. LMP2A does not support ferroptosis resistance.**

(A) Analysis of LMP1 and LMP2A expression in Daudi Burkitt cells. Representative blot of n=3 experiments of whole cell lysates (WCL) from Daudi cells that were mock induced or induced for LMP1 or LMP2A for 24 hours.
(B) Analysis of LMP2A effects on Burkitt cell ferroptosis. Representative FACS analysis of 7-AAD uptake by Daudi cells mock induced or induced for LMP2A and then treated with DMSO, 10µM erastin, 0.1µM ML-210 and/or 2.5µM Fer1 for 48 hours.

**Figure S2. Activation of signaling cascades downstream of TNF, CD40L, or SMAC mimetic**

Model of pathways activated by TNFα, CD40L or SMAC mimetic birinapant. TNFα activates canonical NF-κB and MAP kinase signaling while CD40L stimulation activates canonical and non-canonical NF-κB, and MAP kinases, PI3K, and PKC signaling. SMAC mimetic targets cIAP1/2 for degradation, preventing the proteasomal degradation of NIK and inducing non-canonical NF-κB signaling.

**Figure S3. LMP1 TES2 signaling protects Daudi Burkitt cells independently of canonical NF-κB signaling.**

(A) Validation of NF-κB activation by CD40L, SMAC mimetic and TNFα. Daudi cells were treated for 24 hours with DMSO, Mega CD40L (50 ng/mL), SMAC mimetic birinapant 20μM or TNFα (10ng/mL). WCL was extracted and immunoblots were performed.
(B) Representative FACS %7AAD+ analysis of n=3 independent replicates of Daudi cells stimulated with DMSO control, MegaCD40L (50ng/mL), SMAC mimetic birinapant 20μM, or TNFα (10ng/mL). After 24 hours, cells were treated with DMSO, 10μM erastin and/or 5μM Fer-1. Viability was assessed 48 hours later.
(C) Validation of IKKβ inhibition of canonical NF-κB signaling. Daudi Burkitt cells mock induced or dox induced for expression of LMP1 together with DMSO, 1μM or 5μM IKKβ inhibitor IKK-2 VIII. After 24 hours, WCL was extracted and immunoblot performed.
(D) Representative FACS %7AAD+ analysis of n=3 independent replicates of Daudi cells mock induced or dox induced for expression of LMP1 together with DMSO, 1 μM, or 5 μM IKKβ inhibitor IKKβi. After 24 hours, cells were treated with DMSO, 10 μM erastin and/or 2.5μM Fer-1. Viability was assessed 48 hours later.
(E) Validation of IKKβi inhibition of canonical NF-κB signaling. Akata Burkitt cells mock induced or dox induced for expression of LMP1 together with DMSO, 1μM or 5μM IKKβ inhibitor IKK-2 VIII. After 24 hours, WCL was extracted and immunoblot performed.
(F) Representative FACS %7AAD+ analysis of n=3 independent replicates of Akata cells induced for expression of GFP or WT, TES1m or TES2m LMP1 for 24 hours. Cells were then treated with DMSO, 10μM erastin and/or 2.5μM Fer-1. Viability was assessed 48 hours later.

**Figure S4. TES2 signaling protects Akata and BL-41 Burkitt cells from erastin ferroptosis induction independently of canonical NF-κB signaling.**

(A) Validation of LMP1 mutant expression. Daudi cells were mock induced or dox induced for expression of WT, TES1m or TES2m LMP1. After 24 hours, WCL was extracted and immunoblot performed.
(B) Representative FACS plot of n=3 independent replicates of Daudi cells mock induced or induced for wild-type LMP1 or LMP1 containing point mutations abrogating TES1 or TES2 function for 24 hours, followed by treatment with DMSO, 10 µM erastin, and/or 2.5µM Fer-1 for 48 hours.
(C) Validation of LMP1 mutant expression. BL-41 cells were mock induced or dox induced for expression of WT, TES1m or TES2m LMP1. After 24 hours, WCL was extracted and immunoblot performed using.
(D) Representative percentage of 7-AAD+ cells from n=3 independent replicates of BL-41 cells mock induced or dox induced for WT, TES1m or TES2m LMP1 for 24 hours, followed by treatment with DMSO, 10µM erastin and/or 2.5µM Fer-1 for 48 hours.

P-values were determined by one-sided Fisher’s exact test. * p<0.05, **p<0.005, ***p<0.0005.

**Figure S5. LMP1 protection from erastin ferroptosis is not dependent on TRAF6 or TAK1.**

(A) Validation of TRAF6 knockout. Daudi cells expressing control or TRAF6 targeting sgRNAs for three days were mock-induced or dox induced for WT LMP1. After 24 hours, WCL was extracted and immunoblotted.
(B) FACS %7AAD+ mean + SD values from n=3 independent replicates of Daudi cells expressing control or TRAF6 sgRNA for three days before mock or wild-type LMP1 expression for 24 hours, followed by treatment with DMSO, 10µM erastin and/or 2.5µM Fer-1 for 48 hours.
(C) Validation of TAK1 knockout. Daudi cells expressing control or TAK1 targeting sgRNAs for three days were mock induced or induced for expression of WT LMP1. After 24 hours, WCL was extracted and immunoblotted.
(D) FACS %7AAD+ mean + SD values from n=3 independent replicates of Daudi cells expressing control or TAK1 sgRNA for three days before mock or wild-type LMP1 expression for 24 hours, followed by treatment with DMSO, 10 µM erastin and/or 2.5µM Fer-1 for 48 hours.

**Figure S6. LMP1 protection from erastin ferroptosis is not dependent on TRAF6 or TAK1.**

(A) Representative FACS %7AAD+ analysis of n=3 independent replicates of Daudi cells expressing control or TAK1 sgRNA for three days before mock induction or dox induction of WT LMP1. After 24 hours, cells were treated with DMSO, 10μM erastin and/or 2.5μM Fer-1. Viability was assessed 48 hours later.
(B) Representative FACS %7AAD+ analysis of n=3 independent replicates of Daudi cells expressing control or TRAF6 sgRNA for three days before mock or dox LMP1 induction. After 24 hours, cells were treated with DMSO, 10 μM erastin and/or 2.5μM Fer-1. Viability was assessed 48 hours later.

**Figure S7. Metabolism pathways increased in LMP1 vs TES2m**

MetaboAnalyst metabolite pathway analysis of metabolites significantly upregulated in cells infected by EBV with WT vs TES2m LMP1, as in Figure 4C. Created in BioRender. Burton, E. (2026) <https://BioRender.com/aru2f7k>.

**Figure S8. Exogenous expression of PFKFB4 rescues TES2 mutant LMP1 infected B cell growth**

(A) Normalized PFKFB4 reads from RNAseq analysis of primary human B cells at the indicated days post-infection by the EBV B95.8 strain(68). Shown are mean ± SD values from n=3 replicates.
(B) Assessment of PFKFB4 protein levels during EBV-mediated B cell transformation. Primary circulating B cells were isolated and infected with B95-8 EBV. WCL was extracted at the indicated DPI and immunoblot was performed. DDX1 was used as loading control as it was previously demonstrated not to fluctuate significantly during EBV-mediated transformation(32).
(C) UCSC Genome browser view of LCL *PFKFB4* gene occupancy by the indicated viral EBNA, LMP1-activaed NF-κB transcription factors and histone methylation marks.
(D) Analysis of exogenous PFKFB4 expression effects on WT versus TES2m EBV infected cells. WCL from cells transduced with lentivirus empty vector control or with lentivirus driving stable V5-PFKFB4 expression, as in Figure 5J, was extracted two days post transduction. Immunoblot was performed using the indicated antibodies.

**Figure S9. Validation of PFKFB4 sgRNA knockdown efficiency in Daudi cells**

(A) Representative FACS %7-AAD+ analysis of n=3 independent replicates of GM12878 cells expressing control of PFKFB4 sgRNA for three days followed by treatment with DMSO, 10µM erastin and/or 2.5µM Fer-1 for 48 hours.
(B) Representative FACS %7-AAD+ analysis of n=3 independent replicates of GM15892 cells expressing control of PFKFB4 sgRNA for three days followed by treatment with DMSO, 10µM erastin and/or 2.5µM Fer-1 for 48 hours.
(C) Representative immunoblot of WCL from Cas9+ Daudi cells after three days of control or PFKFB4 sgRNA expression, followed by 24 hours of mock induction or dox induction of LMP1 expression.
(D) Representative FACS %7-AAD+ analysis of n=3 independent replicates of Daudi cells expressing control of PFKFB4 sgRNA for three days followed by mock or LMP1 expression for 24 hours. Cells were then treated with DMSO, 10µM erastin and/or 2.5µM Fer-1 for 48 hours.
