## Supplementary material for "Epstein-Barr Virus Latent Membrane Protein 1 Suppresses Ferroptosis via Pentose Phosphate Pathway and Glutathione Metabolism": Fig. S10

1. **TES2 –(YYD to ID) Mutant** (1685 bps)

agggcccgcctttgatgacagacggaggcGGCGGTCATAGTCATGATTCCggccatggcggcggtgatccacaccttcctacgc (LMP1 sequence upstream- not in gene block)

**tgcttttgggttcttctggttccggtggagatgatgacgacccccacggcccagttcagctaagcATcgaTtaa**cctttctttacttcta cggggaaatgtgcgcggaacccctatttgtttatttttctaaatacattcaaatatgtatccgctcatgagacaataaccctgataaatgcttcaataatattgaaaaaggaagagtATGAGTATTCAACATTTCCGTGTCGCCCTTATTCCCTTTTTTGCGGCATTTTGCCTTCCTGTTTTTGCTCACCCAGAAACGCTGGTGAAAGTAAAAGATGCTGAAGATCAGTTGGGTGCACGAGTGGGTTACATCGAACTGGATCTCAACAGCGGTAAGATCCTTGAGAGTTTTCGCCCCGAAGAACGTTTTCCAATGATGAGCACTTTTAAAGTTCTGCTATGTGGCGCGGTATTATCCCGTATTGACGCCGGGCAAGAGCAACTCGGTCGCCGCATACACTATTCTCAGAATGACTTGGTTGAGTACTCACCAGTCACAGAAAAGCATCTTACGGATGGCATGACAGTAAGAGAATTATGCAGTGCTGCCATAACCATGAGTGATAACACTGCGGCCAACTTACTTCTGACAACGATCGGAGGACCGAAGGAGCTAACCGCTTTTTTGCACAACATGGGGGATCATGTAACTCGCCTTGATCGTTGGGAACCGGAGCTGAATGAAGCCATACCAAACGACGAGCGTGACACCACGATGCCTGTAGCAATGGCAACAACGTTGCGCAAACTATTAACTGGCGAACTACTTACTCTAGCTTCCCGGCAACAATTAATAGACTGGATGGAGGCGGATAAAGTTGCAGGACCACTTCTGCGCTCGGCCCTTCCGGCTGGCTGGTTTATTGCTGATAAATCTGGAGCCGGTGAGCGTGGGTCTCGCGGTATCATTGCAGCACTGGGGCCAGATGGTAAGCCCTCCCGTATCGTAGTTATCTACACGACGGGGAGTCAGGCAACTATGGATGAACGAAATAGACAGATCGCTGAGATAGGTGCCTCACTGATTAAGCATTGGTAAattaccctgttatcccta**ccagttcagctaagcATcgaTtaa**cctttctttacttctaggcattaccatgtcataggcttgcctgactgact

**LMP1 sequence in blue**

**Point Mutations**

16-20bp Short sequence duplication (on both ends of the Amp cassette)

50bp extension sequence

Amp promoter

AmpR cassette

SceI site
